# Expression and distribution of synaptotagmin isoforms in the zebrafish retina

**DOI:** 10.1101/2020.08.06.239814

**Authors:** Diane Henry, Christina Joselevitch, Gary G. Matthews, Lonnie P. Wollmuth

## Abstract

Synaptotagmins belong to a large family of proteins. While various synaptotagmins have been implicated as Ca^2+^ sensors for vesicle replenishment and release at conventional synapses, their roles at retinal ribbon synapses remain incompletely understood. Zebrafish is a widely used experimental model for retinal research. We therefore investigated the homology between human, rat, mouse, and zebrafish synaptotagmins 1 to 10 using a bioinformatics approach. We also characterized the expression and distribution of various synaptotagmin (*syt*) genes in the zebrafish retina using RT-PCR and *in situ* hybridization, focusing on the family members whose products likely underlie Ca^2+^-dependent exocytosis in the central nervous system (synaptotagmins 1, 2, 5 and 7). We find that most zebrafish synaptotagmins are well conserved and can be grouped in the same classes as mammalian synaptotagmins, based on crucial amino acid residues needed for coordinating Ca^2+^ binding and determining phospholipid binding affinity. The only exception is synaptotagmin 1b, which lacks 34 amino acid residues in the C2B domain and is therefore unlikely to bind Ca^2+^ there. Additionally, the products of zebrafish *syt5a* and *syt5b* genes share identity with mammalian class 1 and 5 synaptotagmins. Zebrafish *syt1*, *syt2*, *syt5* and *syt7* paralogues are found in the zebrafish brain, eye, and retina, excepting *syt1b*, which is only present in the brain. The complementary expression pattern of the remaining paralogues in the retina suggests that *syt1a* and *syt5a* may underlie synchronous release and *syt7a* and *syt7b* may mediate asynchronous release or other Ca^2+^ dependent processes in different types of retinal neurons.

## INTRODUCTION

Neuronal communication depends critically on Ca^2+^ (Katz & Miledi, 1965; Miledi, 1973) and Ca^2+^ sensors to regulate vesicle release and replenishment (Neher & Sakaba, 2008). The main Ca^2+^ sensors for exocytosis in the central nervous system of vertebrates are synaptotagmins, a large protein family that comprises about 15-17 members in humans (Sudhof, 2002; Craxton, 2004; Gustavsson & Han, 2009). Eight of these isoforms do not bind Ca^2+^ at all (Dai et al., 2004; Hui et al., 2005). The remainder show distinct Ca^2+^ sensitivities and phospholipid binding properties (Li et al., 1995; Sugita et al., 2002; Bhalla et al., 2005; Hui et al., 2005), as well as varying subcellular distributions (Sugita et al., 2001; Takamori et al., 2006; Dean et al., 2012) and cellular expression (Ullrich et al., 1994), potentially conferring neurons with a highly versatile repertoire in terms of synaptic gain, kinetics and transmission bandwidths (Hui et al., 2005; Xu et al., 2007; Chen & Jonas, 2017). However, only synaptotagmins 1, 2, 5 and 7 have been directly implicated in synaptic transmission (Xu et al., 2007; Bacaj et al., 2013).

In the retina, synapses often rely on continuous transmission. To cope with the demands of uninterrupted sensory inputs, the first retinal neurons - photoreceptors and bipolar cells - use specialized presynaptic contacts called ribbon synapses (Matthews & Fuchs, 2010). Ca^2+^ influx is continuous at such synapses and modulates neurotransmitter release around a mean tonic level. Retinal ribbon synapses presumably depend on multiple modes of release to relay information (Jackman et al., 2009; Oesch & Diamond, 2011), and because photoreceptor and bipolar cell ribbon synapses are distinct morphologically and physiologically (Matthews & Fuchs, 2010), it is plausible that they use different sets of synaptotagmins to control neurotransmitter release.

Although synaptotagmin 1 (Ullrich & Sudhof, 1994; Fox & Sanes, 2007; Grassmeyer et al., 2019), synaptotagmin 2 (Ullrich et al., 1994; Fox & Sanes, 2007; Neumann & Haverkamp, 2013), synaptotagmin 3 (Butz et al., 1999; Berntson & Morgans, 2003) and synaptotagmin 7 (Luo et al., 2015) have been found in the retina, only synaptotagmin 1 (Grassmeyer et al., 2019) and synaptotagmin 7 (Luo et al., 2015) were reported to modulate different aspects of release in retinal ribbon-containing neurons. Furthermore, synaptotagmin 1 is thought to underlie alone both modes of synaptic transmission in mammalian cone but not rod photoreceptors (Grassmeyer et al., 2019). To date, the synaptotagmins controlling transient and sustained release in bipolar cells are unknown.

Zebrafish are a powerful animal model to study the nervous system in general and the retina specifically. First, there are multiple available toolboxes for the manipulation of genes of interest (Kawakami, 2007; Ablain et al., 2015; Kawakami et al., 2016; Niklaus & Neuhauss, 2017; Wierson et al., 2020). Second, zebrafish husbandry is relatively easy, mating has high yields and development is fast when compared to mammalian models. Lastly, zebrafish are experimentally accessible during development and display conserved retinal structures and main cell types (Gestri et al., 2012; Angueyra & Kindt, 2018).

The validity of this animal model for the study of retinal structure and function depends critically on the ability to draw analogies between the retinal anatomy, physiology and molecular biology of zebrafish and mammals. We therefore investigated the homology, expression, and distribution of synaptotagmins in the zebrafish retina. To do so, we took advantage of bioinformatic approaches to identify the genetic, sequence, and potential structural homology between zebrafish, mouse, and human synaptotagmins. We also assayed for the presence of synaptotagmins in the retina using RT-PCR and *in situ* hybridization.

We find that zebrafish synaptotagmins are well conserved and can be grouped in the same classes as mammalian synaptotagmins. The only exception are synaptotagmins 5a and 5b, which are at an intermediate position between mammalian class 1 and 5 synaptotagmins.

Zebrafish proteins retain key amino acid residues needed for coordinating Ca^2+^ binding and determining phospholipid binding affinity. Notably, while zebrafish synaptotagmin 1a is homologous to mammalian synaptotagmin 1 and is expressed in the retina, synaptotagmin 1b is not, and lacks the residues necessary to function as a Ca^2+^ sensor in its C2B domain. Zebrafish *syt1*, *syt2*, *syt5* and *syt7* paralogues are found in the zebrafish brain, eye and retina, except *syt1b*, which is only present in the brain. The remaining paralogues had complementary retinal expression patterns, suggesting that *syt1a* and *syt5a* may underlie synchronous release and *syt7a* and *syt7b* may mediate asynchronous release in different types of retinal neurons.

## MATERIALS AND METHODS

### Animals

All animal procedures were approved by the institutional animal care and usage committee (IACUC) at Stony Brook University and were in concordance with the guidelines established by the National Institutes of Health and by the Statement for the Use of Animals in Ophthalmic and Vision Research from The Association for Research in Vision and Ophthalmology.

Wild-type (WT) zebrafish (*Danio rerio*) were kept at 28.5 °C in aquaria under a 13:11 h light to dark cycle and fed artemia and GEMMA micropellets twice a day. The WT strain used for all experiments was a hybrid WT background consisting of Tubingen long-fin crossed to Brian’s WT. Adult animals (age 8-12 months) of either sex were dark-adapted for 2 h and euthanized by immersion in 1 mM tricaine methanesulfonate (SIGMA, cat. no. A5040) in pH 7.0 buffered system water prior to experiments.

### RT-PCR analysis

Zebrafish eyes and brains were dissected from adult animals and rapidly frozen by immersion in a mixture of ethanol and dry ice. Total RNA was extracted by the method of Cathala *et al.* (Cathala et al., 1983) and stored in ethanol at −80 °C until use.

Reverse transcription using Superscript IV reverse transcriptase (Invitrogen) was performed according to the manufacturer’s protocol. In brief, 250-500 μg of total DNAse I-treated total RNA was added to nuclease-free water to a final volume of 11 μL. Subsequently, 1 μL of random hexamers and 1 μL of 10 mM dNTP mix were added to the RNA solution, heated to 65 °C for 5 minutes and then incubated on ice for one minute. After addition of 4 μL of 5x buffer, 1 μL 10 mM DTT and 1 μL RNase inhibitor, the sample was incubated at 23°C and 1 μL of Superscript IV reverse transcriptase was added; synthesis was performed for 1 hour at 55 °C. The enzyme was inactivated at 80 °C for 10 minutes. All reagents were obtained from Invitrogen/Thermo Fisher.

For conventional PCR, 1 μL reverse-transcribed cDNA was added to 25 μL of PCR buffer and reactants, using EconoTaq DNA polymerase (Lucingen). The standard amplification protocol consisted of 95°C for 5 minutes, followed by 35 cycles of 95, 55 and 72°C for 1 minute each, and ending with a 72°C-extension for 5 minutes. PCR primers were designed based on GenBank sequences for zebrafish synaptotagmin isotypes. Type-specific primers were designed such that each one amplified a unique fragment either near the ATG start of the respective coding sequence (CDS) or a larger unique fragment in the untranslated (UTR) region. The design of isotype-specific primers for synaptotagmin genes is complicated by the fact that nearly 40% of the gene is comprised of homologous C2A and C2B domains. The primers shown in **Table 1** were designed to amplify the largest unique fragment for each gene. DNA fragments were confirmed by sequencing and alignment. cDNA products of correct sizes were gel-purified and subcloned into the pGEM-T Easy cloning vector (Promega). Selected clones were sequenced on an automatic DNA sequencer and aligned to GenBank (https://www.ncbi.nlm.nih.gov/genbank/, RRID:SCR_002760) data to verify the integrity of DNA fragments.

**Table 1.**
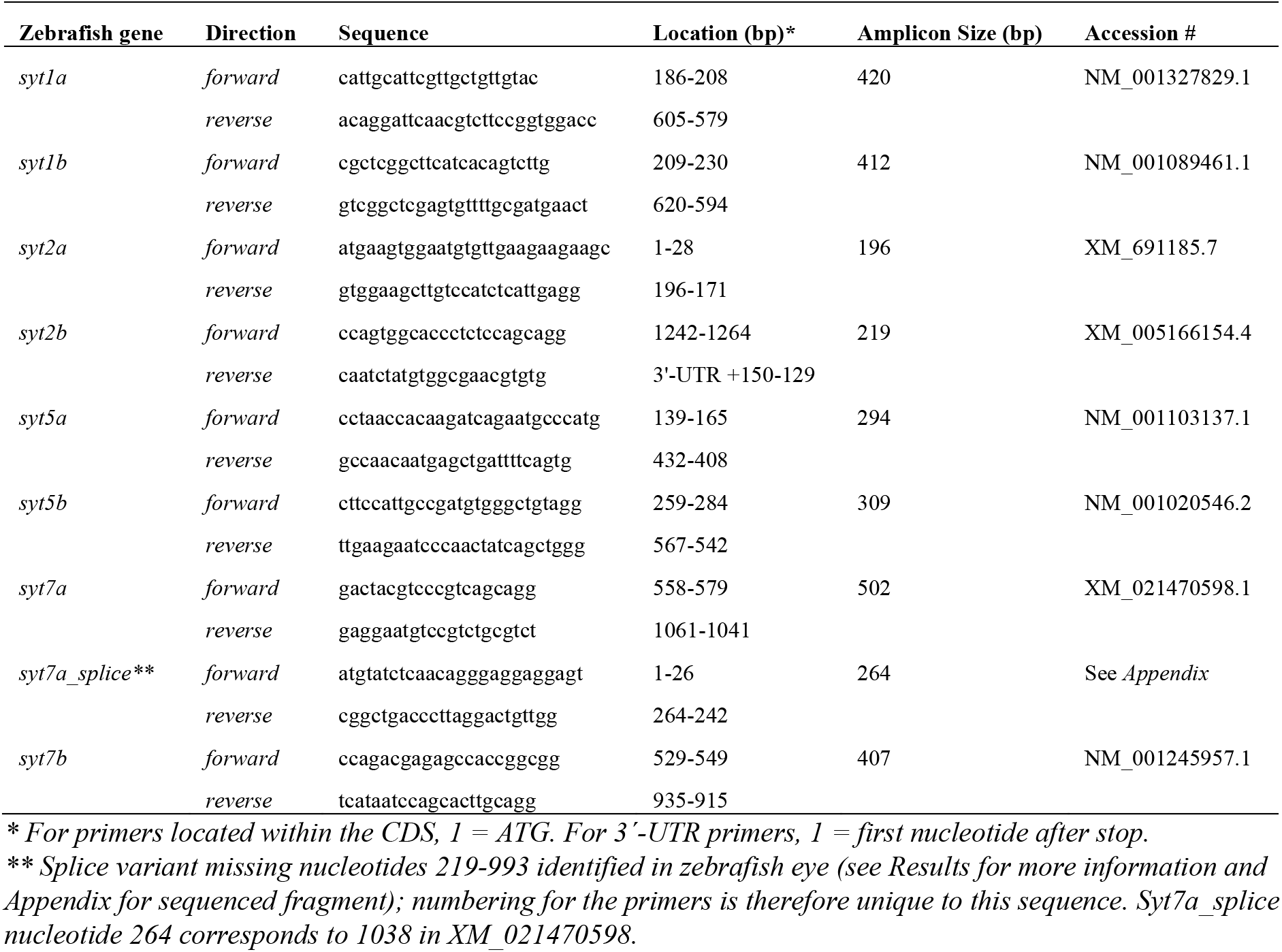
PCR primers.

### *In situ* hybridization

Zebrafish eyes were dissected and fixed in 4% paraformaldehyde in phosphate-buffered saline (PBS) overnight at 4 °C. The fixed tissue was washed in PBS and cryoprotected in 30% sucrose before embedding in Shandon M1 media (Thermo Fisher Scientific) and stored at −80 °C until use. On day one of *in situ* experiment, 25-μm sections were placed on Superfrost Plus microscope slides (Thermo Fisher Scientific) and dried at 55°C for a minimum of 3 hours. This tissue was then immediately used for *in situ* hybridization.

Isotype-specific probes for zebrafish synaptotagmins present in the eye were synthesized in sense and antisense directions from cDNA obtained by RT-PCR. Riboprobe lengths and locations are listed in **Table 2**. All probes were localized in the 5’ end of the gene or 3’-UTR, where sequences are divergent. Synthesis, hybridization, and detection of digoxigenin-labeled probes were carried out according to manufacturer’s protocol (Roche), using anti-digoxigenin antibody conjugated to alkaline phosphatase for detection. A SNAP25b riboprobe (a gift from H. Sirotkin, Moravec et al., 2016) was used as a positive control for tissue and reagents.

**Table 2.**
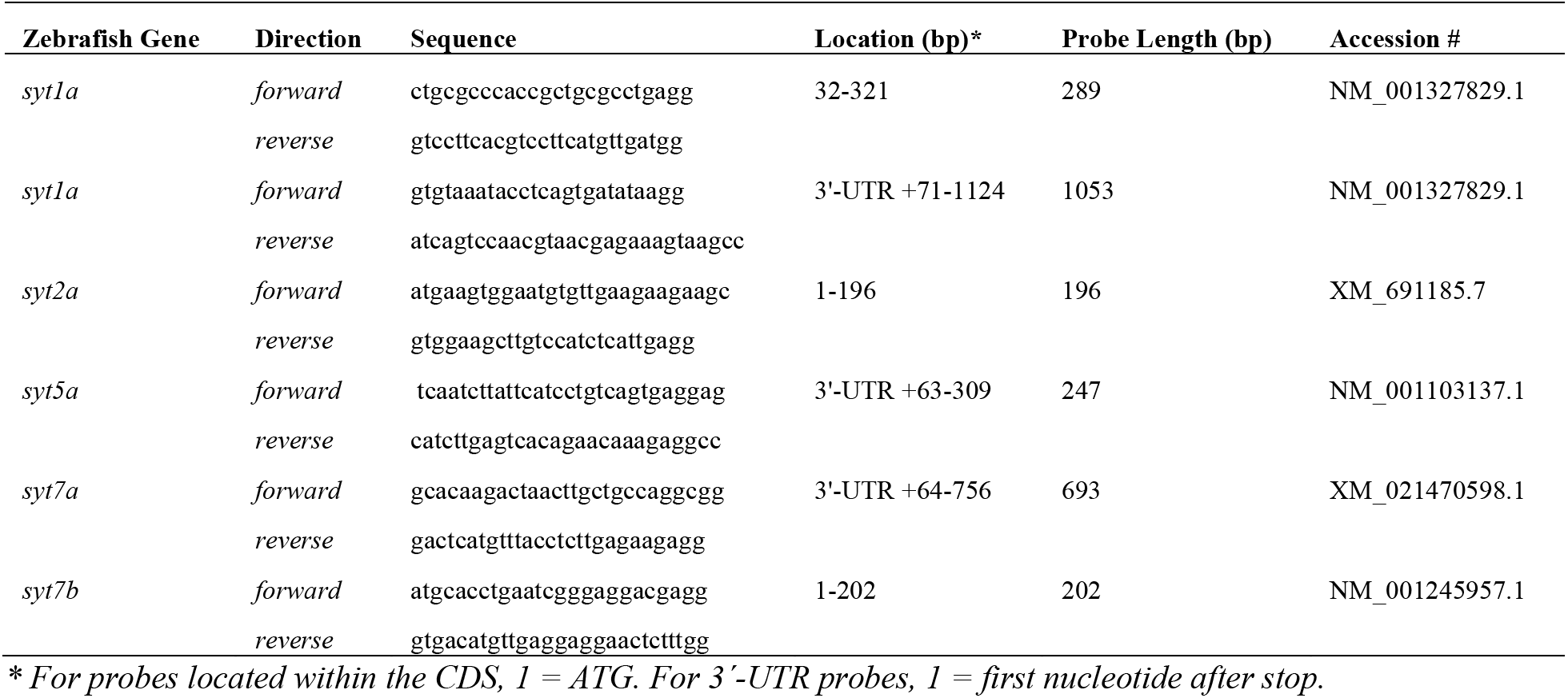
Riboprobes for *in situ hybridization*.

Slides were viewed and photographed under 20x or 40 x magnification (Zeiss PlanNeofluor objectives, 0.5 and 0.75 N.A., respectively) in a microscope equipped for brightfield (Zeiss Axioskop) attached to a color camera (Infinity 3, Lumenera Corporation, Ottawa, Canada) controlled by Infinity Capture software (Lumenera Corporation, Ottawa, Canada). Photomicrographs of sense and anti-sense pairs generated from the same experiment were acquired as TIFF files at 1936×1456 pixels and post-processed jointly with Adobe Photoshop (https://www.adobe.com/products/photoshop.html, RRID:SCR_014199) to correct contrast and brightness.

### Bioinformatics

The orthology between the zebrafish *syt* genes and their human and mouse counterparts was determined by searching the Zebrafish Information Network (Zfin, http://zfin.org, RRID:SCR_002560), NCBI Entrez Gene (http://www.ncbi.nlm.nih.gov/gene, RRID:SCR_002473), NCBI Homologene (http://www.ncbi.nlm.nih.gov/homologene, RRID:SCR_002924) and Ensembl (http://www.ensembl.org/, RRID:SCR_002344) databases. Additionally, the NCBI BLAST Suite (https://blast.ncbi.nlm.nih.gov/Blast.cgi, RRID:SCR_004870) was used to align and compare gene, transcript or protein sequences of interest and to investigate possible interactors with *in situ* probes or help determine homology.

In addition to percent identity, we also used two scores in Ensembl to evaluate homology: Gene Order Conservation (GOC) and the Whole Genome Alignment (WGA) scores. The GOC score, which ranges from 0 to 100, indicates how many of the four closest neighbors of a gene match between orthologous pairs, and assumes that genes that are descended from the same gene are likely to be part of a block of genes, all in the same order, in both species. The WGA score also varies from 0 to 100 and indicates how well query and target genomic regions align to each other It assumes that genes which are orthologous to each other will fall within genomic regions that can be aligned to one another (i.e. percent synteny).

For phylogenetic analysis, protein sequences from NCBI Protein (http://www.ncbi.nlm.nih.gov/protein, RRID:SCR_014312) databases were aligned with Clustal Omega (https://www.ebi.ac.uk/Tools/msa/clustalo/, RRID:SCR_001591, Sievers et al., 2011) or MUSCLE (https://www.ebi.ac.uk/Tools/msa/muscle/, RRID:SCR_011812, Edgar, 2004), both at default settings for highest accuracy. After alignment, regions with gaps or misaligned were removed with GBlocks (http://molevol.cmima.csic.es/castresana/Gblocks_server.html, RRID:SCR_015945, Castresana, 2000) to eliminate ambiguity. A stringent set of parameters was chosen to eliminate divergent regions and improve reliability: (i) no gaps allowed; (ii) blocks after gap cleaning ≥ 10 residues; (iii) contiguous nonconserved residues ≤ 4; (iv) sequences for conserved position ≥ 28; and (v) sequences for a flanking position ≥ 45. After curation, 117 amino acids were selected for subsequent evaluation.

Results were imported into JalView (https://www.jalview.org/, RRID:SCR_006459, Waterhouse et al., 2009) for visualization and calculation of neighbor joining phylogenetic trees using the BLOSUM62 matrix. Phylograms were imported as Newick files into iTOL (Interactive Tree of Life, https://itol.embl.de, RRID:SCR_004473, Ciccarelli et al., 2006; Letunic & Bork, 2019) for manipulation and annotation, and exported as vector-based graphics (i.e. svg or eps files) for final editing in Canvas X (http://www.canvasgfx.com/en/products/canvas-15, RRID:SCR_014312).

Prediction of secondary structure and 3-dimensional modeling of zebrafish synaptotagmin sequences was performed with RaptorX (http://raptorx.uchicago.edu/, RRID:SCR_018118). Models were visualized and aligned to known crystal structures of mammalian synaptotagmins obtained from the Research Collaboratory for Structural Bioinformatics Protein Data Bank (RCSB PDB, http://www.rcsb.org/pdb/, RRID:SCR_012820) using PyMOL (http://www.pymol.org/, RRID:SCR_000305) and exported as PNG files at 5000 x 5000 pixel resolution for subsequent editing in Canvas X.

## RESULTS

### Homology of zebrafish synaptotagmins to their mammalian counterparts

Due to a whole genome duplication that happened at the beginning of the teleost lineage some 320-350 million years ago (Amores et al., 1998; Glasauer & Neuhauss, 2014; Pasquier et al., 2016), zebrafish in general have two synaptotagmin (*syt*) paralogue genes, named *a* and *b*, for each mammalian counterpart. The exceptions are *syt3*, *syt4*, *syt8* and *syt10* (Craxton, 2004, 2010), which have only one gene, probably as a result of deleterious mutations of one of the paralogues that led to loss of function (Postlethwait et al., 2000; Glasauer & Neuhauss, 2014; Pasquier et al., 2016). At present it is unknown whether the remaining zebrafish *syt* genes still act as Ca^2+^ sensors, or whether they have acquired new functions during evolution.

To begin to address this issue, we compiled the genomic identities between zebrafish and human orthologues in the Ensembl database (**Figure 1A**) and compared them to those between mouse and human orthologue pairs (**Figure 1B**). For this and subsequent analyses, we concentrated on synaptotagmins 1 to 10, since synaptotagmins 11-15 do not bind Ca^2+^ (von Poser et al., 1997; Fukuda, 2003a; Bhalla et al., 2008; Craxton, 2010), and there is uncertainty as to whether synaptotagmins 16-17 should be considered members of the synaptotagmin family because they lack a transmembrane domain (Gustavsson & Han, 2009; Wolfes & Dean, 2020). The nomenclature between zebrafish and mammalian orthologue pairs is consistent (i.e., the human orthologue for zebrafish syt1a is human SYT1; Craxton, 2004), because zebrafish *syt* genes were named after extensive examination of their syntenic relations and the amino acid sequences of their products (Craxton, 2004, 2010).

**Figure 1.**
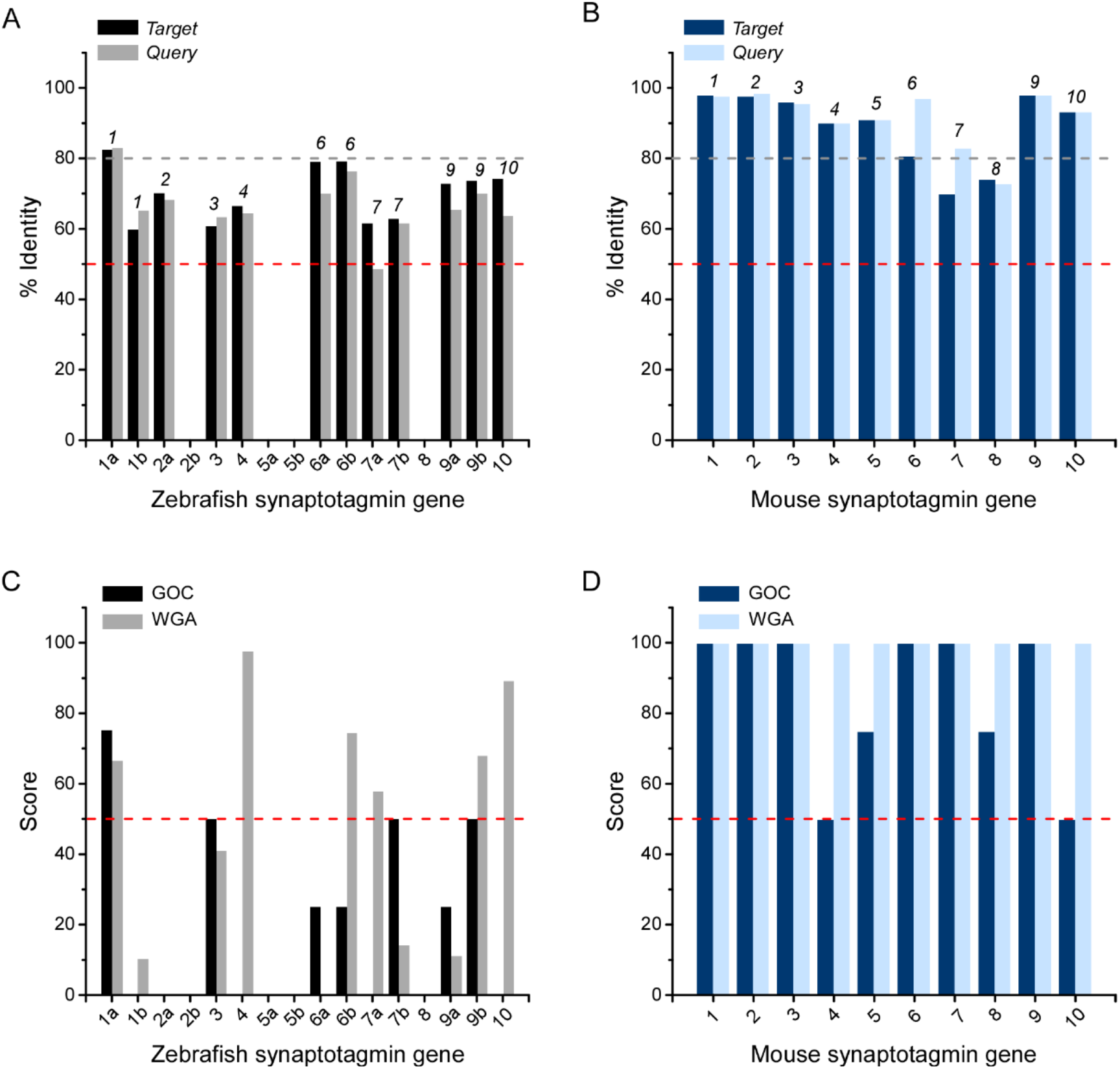
Genomic homology between zebrafish, mouse and human synaptotagmin 1 to 10. (A, B) The percent identity listed in the Ensembl database between zebrafish and human (A) and between mouse and human (B) orthologues (human *SYT* gene numbers are indicated above the bars). *Target*: % of the human sequence matching the zebrafish or mouse sequence; *Query*: % of the zebrafish or mouse sequence matching the human sequence. (A) The percent identity between zebrafish and human is above 50% (dashed red line), except for *syt7a* (*query*: 62%; *target:* 49%). The only zebrafish gene with >80% identity to its human counterpart (dashed grey line) is *syt1a* (*query*: 82.5%; *target:* 83%). Empty spaces in the graph reflect that zebrafish *syt2b* and *syt8* are not mapped in Ensembl, and that the database does not list any human orthologue gene matches for zebrafish *syt5a* and *syt5b*. Zebrafish has duplicate genes for all synaptotagmins, except for *syt*3, *syt*4, *syt*8 and *syt10*. (B) The percent identity between mouse and human is in most cases above 80% (dashed grey line), except for *Syt7* (*query*: 70%; *target:* 83%) and *Syt8* (*query*: 74%; *target:* 73%). (C, D) Gene Order Conservation (*GOC)* and Whole Genome Alignment (*WGA)* scores, which vary from 0 to 100, are an additional means to evaluate homology (see *Materials & Methods*). Both scores are much lower for the zebrafish-human (C) than for mouse-human (D) orthologues.

Although the percent identity between zebrafish and human orthologues (**Figure 1A**) is lower than that between mouse and human counterparts (**Figure 1B**), in almost all cases identity is >50%, the only exception being *syt7a*. GOC and WGA scores were much lower for zebrafish-human pairs (**Figure 1C**) than those between mouse and human orthologue genes (**Figure 1D**).

Some gene pairs are missing in **Figure 1A** and **Figure 1C**: zebrafish *syt2b* and *syt8* genes have not been mapped in Ensembl, and although *syt5a* and *syt5b* are mapped, human or murine orthologue searches for these paralogues did not yield any hits in either Ensembl or NCBI. Indeed, if these genes are true orthologues as suggested (Craxton, 2004, 2010), they must have diverged considerably, since the amino acid similarity between human synaptotagmin 5 and zebrafish synaptotagmins 5a and 5b is low (between 49 and 51% identity for synaptotagmin 5a, depending on the isoforms compared, and 53% for synaptotagmin 5b). Aligning the translation products of the *syt5a* and *syt5b* paralogues with those of human SYT genes with MUSCLE yielded human synaptotagmin 2 as the closest match for zebrafish synaptotagmin 5a (85% and 61% identity, respectively), and human synaptotagmin 1 as the closest match for zebrafish synaptotagmin 5b (61% identity) and synaptotagmin 8 (between 53 and 59% identity, depending on the isoforms compared). That said, the manually curated orthology from the Zfin database indicates human *SYT5* and murine *Syt5* as the orthologues for *syt5a* and *syt5b* based on both synteny and amino acid sequence, albeit with no quantitative measure of homology.

Presumably, such sequence divergence arises because of the accumulation of mutations over many generations, which may or may not impair protein function (Glasauer & Neuhauss, 2014). On the other hand, one expects functional portions of the translation products to be more conserved amongst different species than those that are not crucial for protein function.

Synaptotagmins are a fairly conserved family of proteins that share a common structure (**Figure 2A**), consisting of a short extracellular/intravesicular N-terminus, a transmembrane domain, and an intracellular portion formed by an intracellular linker, two tandemly arranged C2 domains interspersed by a shorter linker sequence, and a C-terminus (Sudhof, 2002; Wolfes & Dean, 2020). The C2 domains, named C2A and C2B, are responsible for Ca^2+^, phospholipid and SNARE binding, and are together the main functional components of these proteins.

**Figure 2.**
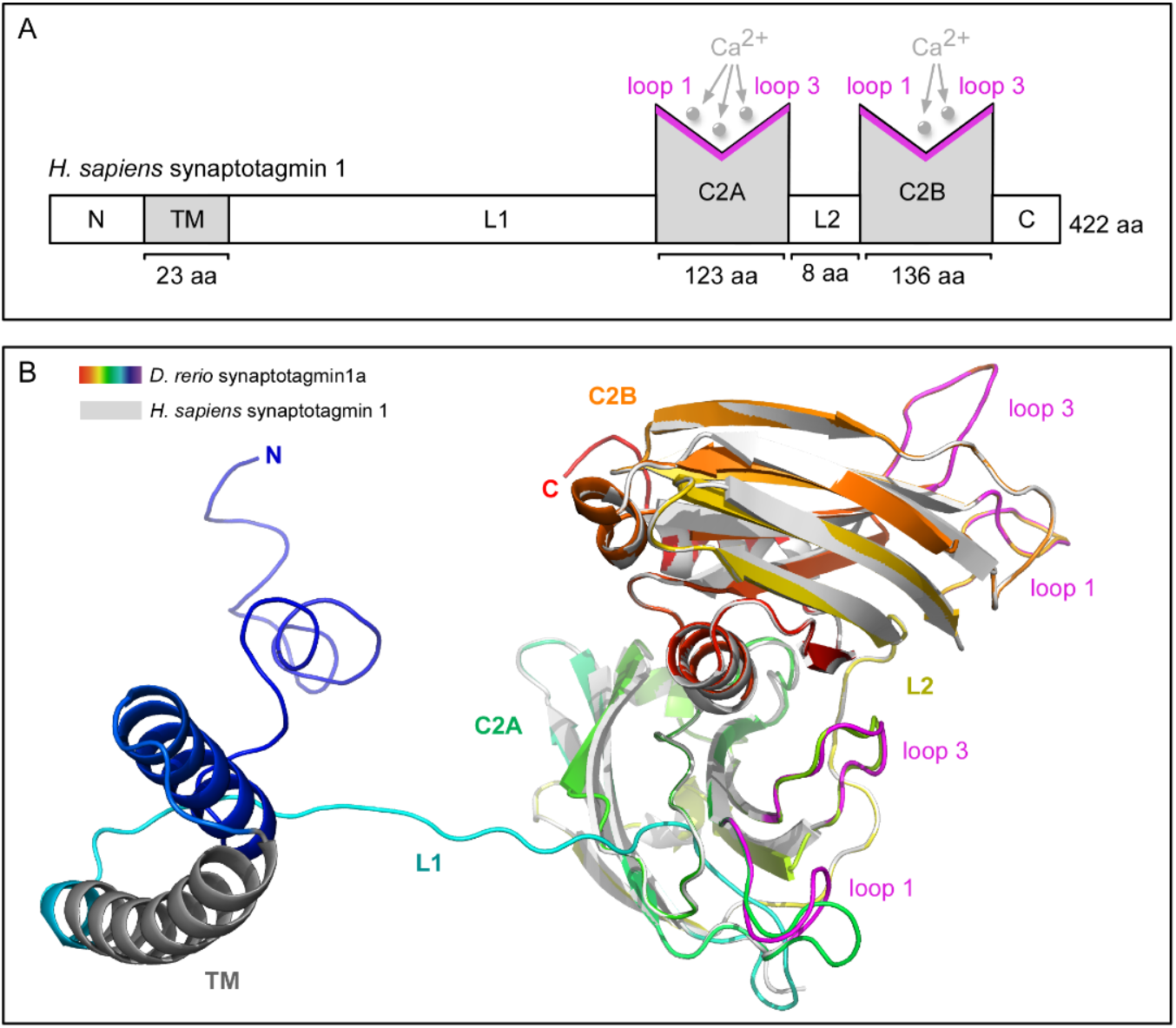
General synaptotagmin structure. (A) Schematic drawing of human synaptotagmin 1. All synaptotagmins share the same basic structure: an intravesicular or extracellular N-terminus (*N*) of variable length; a transmembrane domain (*TM*) that anchors the protein to the vesicular or plasma membrane; a variable linker region (*L1*); two almost identical Ca^2+^-binding domains (*C2A* and *C2B*) arranged in tandem and coupled by a second shorter linker sequence (*L*2); and a C-terminus (*C*). The number of amino acid residues of the non-variable regions for human synaptotagmin 1 are shown. The loops responsible for Ca^2+^ binding in the C2 domains (loops 1 and 3) are shown in pink. Drawing not to scale. (B) Ribbon model of the 3-dimensional structure of zebrafish synaptotagmin 1a (rainbow colors) generated with RaptorX from the amino acid sequence (isoform 1, accession number in **Table 3**) and superposed on the crystal structure of the C2 domains of human synaptotagmin 1 (shown in grey, PDB: 2R83; Fuson et al., 2007). Loops responsible for Ca^2+^ binding in the C2 domains (loops 1 and 3) of the human synaptotagmin are shown in pink.

The modeled 3-D structure of the C2 domains of zebrafish synaptotagmin 1a (depicted in rainbow colors in **Figure 2B**) align to the crystal structure of human synaptotagmin 1 (in grey, **Figure 2B**) (PDB: 2R83, Fuson et al., 2007). This is highly suggestive of functional conservation during evolution. To look more closely at the adaptive radiation of the functional domains of zebrafish synaptotagmins, we examined the phylogenetic relations of conserved amino acid sequences in the C2 domains of zebrafish and human synaptotagmins 1 to 10 (**Figure 3**). The NCBI accession #s used for this analysis are compiled in **Table 3** and **Table 4;** sequences from NCBI were chosen because Ensembl is incomplete (i.e. no zebrafish *syt2b* or *syt8* genes and products).

**Table 3.**
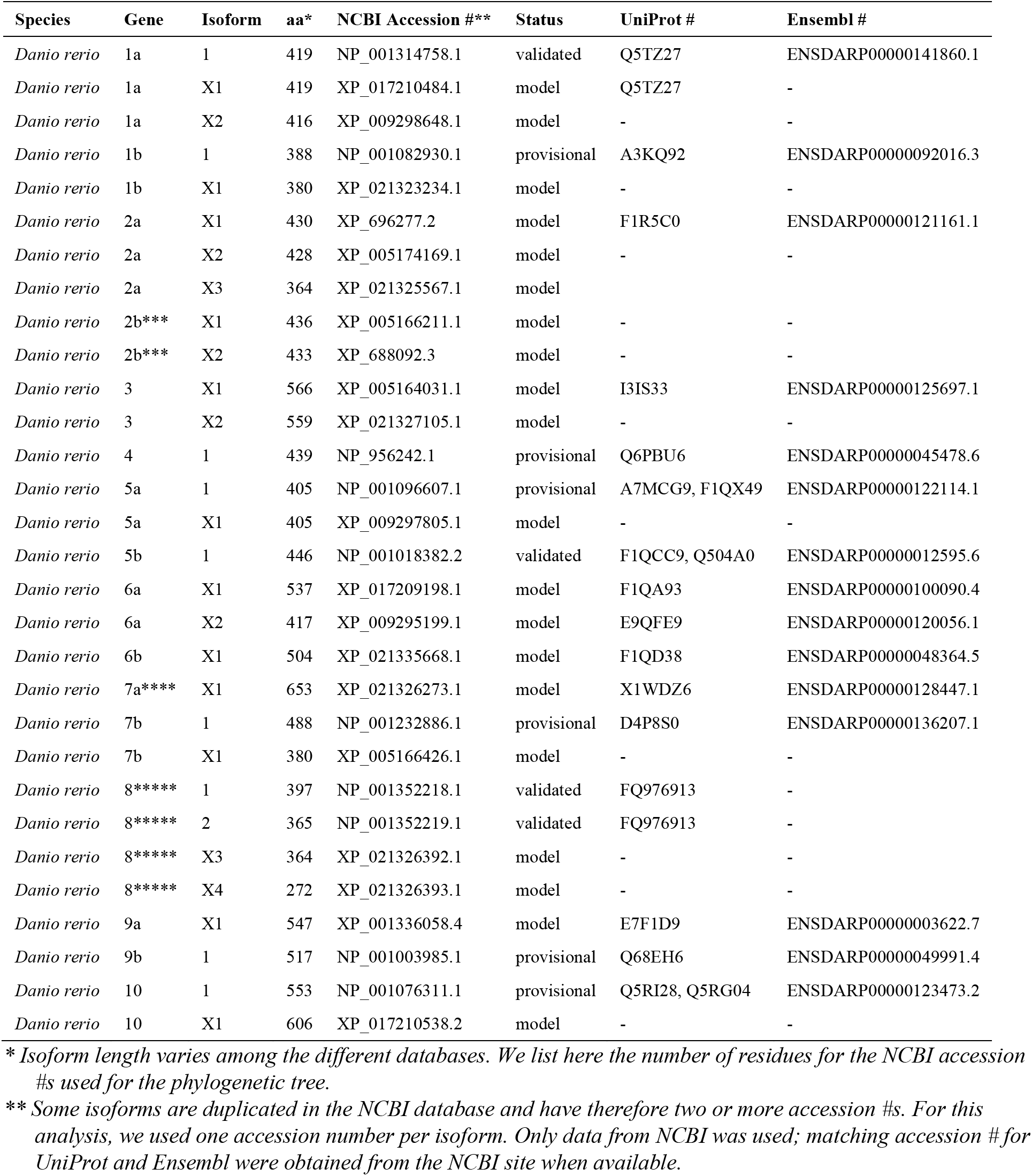
Accession data for the zebrafish synaptotagmins used for phylogenetic analysis.

| Species | Gene | Isoform | aa* | NCBI Accession #** | Status | UniProt # | Ensembl # |
| --- | --- | --- | --- | --- | --- | --- | --- |
| *Danio rerio* | 1a | 1 | 419 | NP_001314758.1 | validated | Q5TZ27 | ENSDARP00000141860.1 |
| *Danio rerio* | 1a | X1 | 419 | XP_017210484.1 | model | Q5TZ27 | - |
| *Danio rerio* | 1a | X2 | 416 | XP_009298648.1 | model | - | - |
| *Danio rerio* | 1b | 1 | 388 | NP_001082930.1 | provisional | A3KQ92 | ENSDARP00000092016.3 |
| *Danio rerio* | 1b | X1 | 380 | XP_021323234.1 | model | - | - |
| *Danio rerio* | 2a | X1 | 430 | XP_696277.2 | model | F1R5C0 | ENSDARP00000121161.1 |
| *Danio rerio* | 2a | X2 | 428 | XP_005174169.1 | model | - | - |
| *Danio rerio* | 2a | X3 | 364 | XP_021325567.1 | model | - | - |
| *Danio rerio* | 2b*** | X1 | 436 | XP_005166211.1 | model | - | - |
| *Danio rerio* | 2b*** | X2 | 433 | XP_688092.3 | model | - | - |
| *Danio rerio* | 3 | X1 | 566 | XP_005164031.1 | model | I3IS33 | ENSDARP00000125697.1 |
| *Danio rerio* | 3 | X2 | 559 | XP_021327105.1 | model | - | - |
| *Danio rerio* | 4 | 1 | 439 | NP_956242.1 | provisional | Q6PBU6 | ENSDARP00000045478.6 |
| *Danio rerio* | 5a | 1 | 405 | NP_001096607.1 | provisional | A7MCG9, F1QX49 | ENSDARP00000122114.1 |
| *Danio rerio* | 5a | X1 | 405 | XP_009297805.1 | model | - | - |
| *Danio rerio* | 5b | 1 | 446 | NP_001018382.2 | validated | F1QCC9, Q504A0 | ENSDARP00000012595.6 |
| *Danio rerio* | 6a | X1 | 537 | XP_017209198.1 | model | F1QA93 | ENSDARP00000100090.4 |
| *Danio rerio* | 6a | X2 | 417 | XP_009295199.1 | model | E9QFE9 | ENSDARP00000120056.1 |
| *Danio rerio* | 6b | X1 | 504 | XP_021335668.1 | model | F1QD38 | ENSDARP00000048364.5 |
| *Danio rerio* | 7a**** | X1 | 653 | XP_021326273.1 | model | X1WDZ6 | ENSDARP00000128447.1 |
| *Danio rerio* | 7b | 1 | 488 | NP_001232886.1 | provisional | D4P8S0 | ENSDARP00000136207.1 |
| *Danio rerio* | 7b | X1 | 380 | XP_005166426.1 | model | - | - |
| *Danio rerio* | 8***** | 1 | 397 | NP_001352218.1 | validated | FQ976913 | - |
| *Danio rerio* | 8***** | 2 | 365 | NP_001352219.1 | validated | FQ976913 | - |
| *Danio rerio* | 8***** | X3 | 364 | XP_021326392.1 | model | - | - |
| *Danio rerio* | 8***** | X4 | 272 | XP_021326393.1 | model | - | - |
| *Danio rerio* | 9a | X1 | 547 | XP_001336058.4 | model | E7F1D9 | ENSDARP00000003622.7 |
| *Danio rerio* | 9b | 1 | 517 | NP_001003985.1 | provisional | Q68EH6 | ENSDARP00000049991.4 |
| *Danio rerio* | 10 | 1 | 553 | NP_001076311.1 | provisional | Q5RI28, Q5RG04 | ENSDARP00000123473.2 |
| *Danio rerio* | 10 | X1 | 606 | XP_017210538.2 | model | - | - |

\* Isoform length varies among the different databases. We list here the number of residues for the NCBI accession #s used for the phylogenetic tree.
\*\* Some isoforms are duplicated in the NCBI database and have therefore two or more accession #s. For this analysis, we used one accession number per isoform. Only data from NCBI was used; matching accession # for UniProt and Ensembl were obtained from the NCBI site when available.
\*\*\* Called synaptotagmin-2-like (LOC559637) in the NCBI database, but synaptotagmin 2b at Zfin, after manual curation (Craxton, 2004).
\*\*\*\* Synaptotagmin 7a has splice variants not listed in NCBI database. This is the longest version (syt7a\_202 in Ensembl).
\*\*\*\*\* Synaptotagmin 8 was formerly called synaptotagmin-1-like.

**Table 4.**
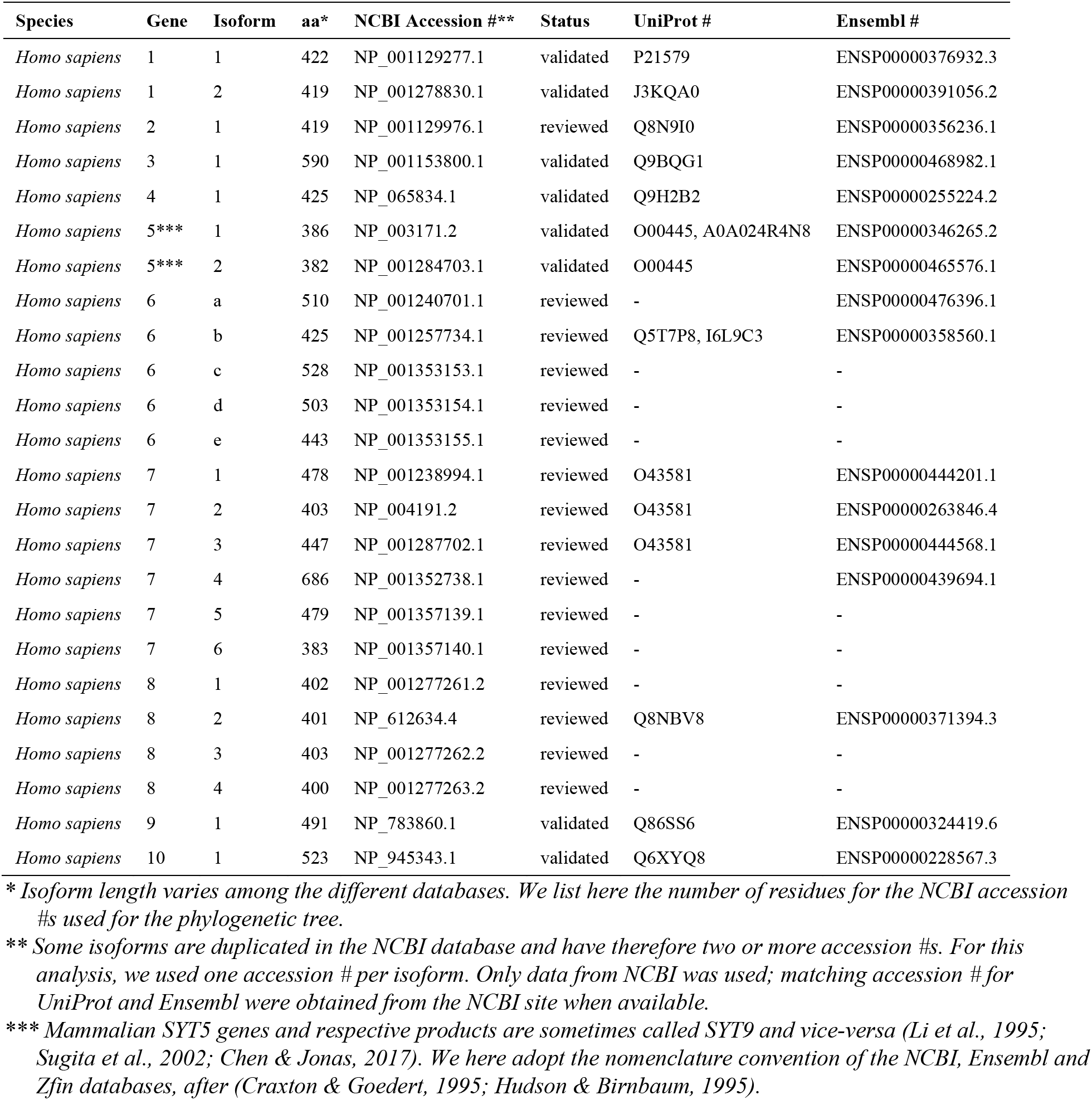
Accession data for the human synaptotagmins used for phylogenetic analysis.

**Figure 3.**
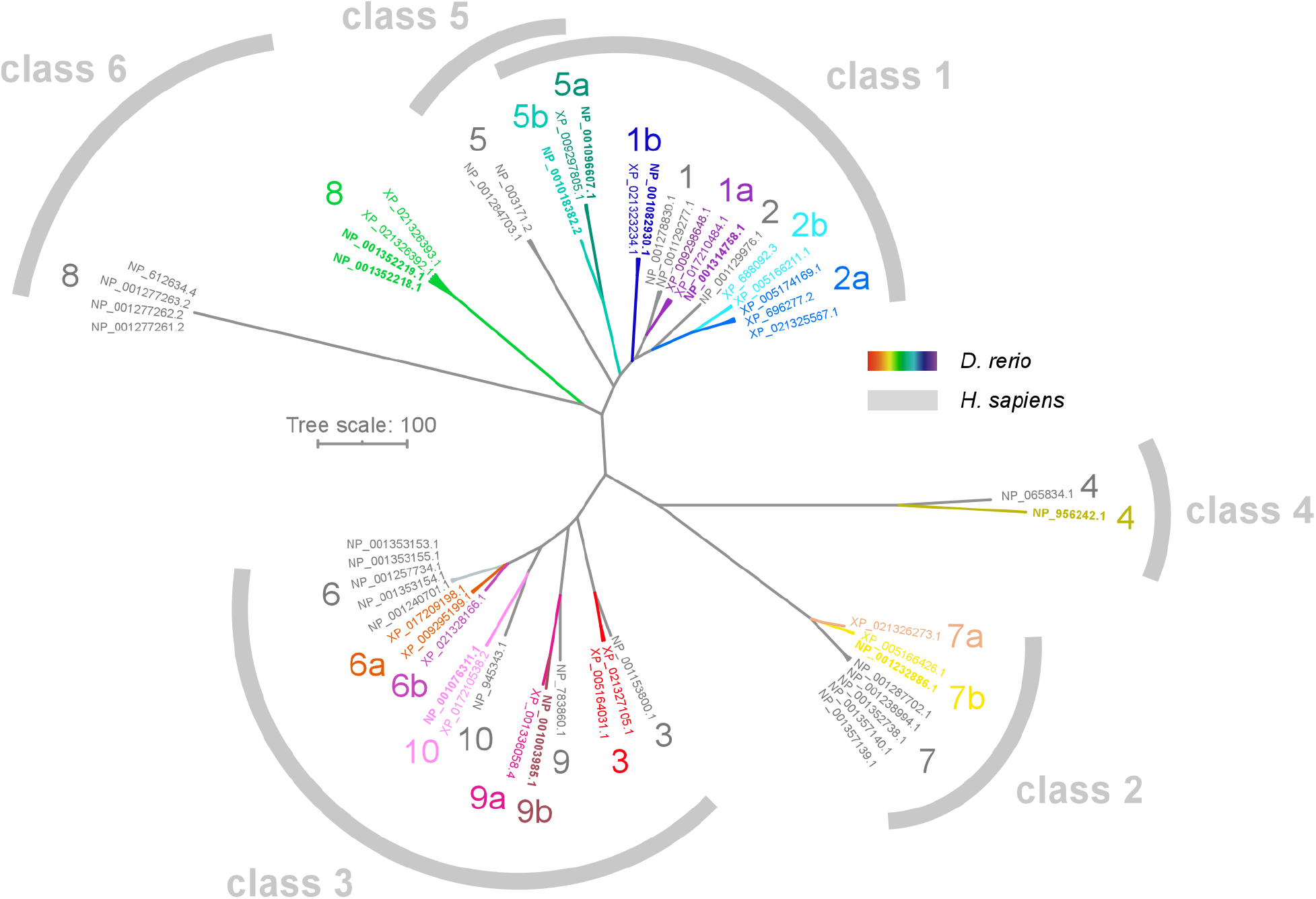
Phylogenetic comparison of zebrafish and human synaptotagmins. Unrooted phylogram of zebrafish and human synaptotagmins 1 to 10 based on 117s conserved amino acid residues from the C2 domains. Proteins are either shown in color (zebrafish) or in grey (human). Class division indicated by brackets, after (Sudhof, 2002). NCBI accession numbers are shown at the end of the branches and represent different isoforms of each protein. Numbers starting with “NP” are validated sequences (shown in bold for zebrafish proteins, see **Table 3** and **Table 4** for NCBI status of each sequence), while numbers starting with “XP” are sequences predicted by computer algorithms from mRNA or genomic sequences, which still await validation. Only validated or reviewed human sequences were used, while for zebrafish, all isoforms listed at the NCBI database were used due to the limited number of validated sequences available. Most zebrafish synaptotagmins fall within the corresponding mammalian class, except synaptotagmins 5a and 5b. These occupy an intermediate location between class 5 and class 1 synaptotagmins (overlapping brackets). Scale bar corresponds to amino acid similarity.

Zebrafish and human orthologue proteins cluster in the same general classes (**Figure 3**) (Sugita et al., 2002). Synaptotagmins 5a and 5b constitute the only exceptions because they fall between class 1 and 5 and could therefore belong to either (overlapping brackets in **Figure 3**). The properties of these classes in mammals and relevant literature are summarized in **Table 5**. Class 1 proteins (synaptotagmins 1 and 2) are fast, low sensitivity and low affinity Ca^2+^ sensors located in vesicles and involved in synchronous synaptic transmission. Class 2 proteins (synaptotagmin 7) are also involved in synaptic transmission but anchor to the plasma membrane and have higher affinity and sensitivity to Ca^2+^ than class 1 synaptotagmins. Class 3 proteins (synaptotagmins 3, 6, 9 and 10) are mostly anchored to the plasma membrane, and regulate endocytosis, postsynaptic receptor trafficking and non-synaptic secretion. Class 4 proteins (synaptotagmins 4 and 11) are insensitive to Ca^2+^ and modulate the activity of other synaptotagmins. Class 5 proteins (synaptotagmin 5) are also fast, low sensitivity/low affinity Ca^2+^ sensors in mammals, but with slower kinetics than class 1 proteins. Finally, class 6 proteins (synaptotagmin 8) are also unable to bind Ca^2+^ and act as inhibitory proteins, as class 4 synaptotagmins.

**Table 5.** Characteristics of neuronal synaptotagmins 1 to 10 in mammals.

| Class* | Syt | S<sub>(Ca)</sub>** | PL_b(Ca)*** | Brain<br>Elsewhere**** | Location | Target | Process | Description |
| --- | --- | --- | --- | --- | --- | --- | --- | --- |
| 1 | 1 | low<sup>1,2</sup> | yes<sup>3,4</sup> | 14.8 | synapses<sup>5</sup> | synaptic vesicles<sup>5,6</sup> | exocytosis<sup>7,8</sup> | synchronous release<sup>9,10</sup> |
| 1 | 2 | low<sup>2</sup> | yes<sup>3,4</sup> | 1.1 | synapses<sup>5</sup> | synaptic vesicles<sup>5</sup> | exocytosis<sup>10</sup> | synchronous release<sup>10</sup> |
| 2 | 7 | high<sup>1,2</sup> | yes<sup>3,4</sup> | 1.1 | synapses<sup>11</sup>, axons<sup>5</sup> | plasma membrane<sup>11</sup> | exocytosis<sup>9,12</sup> | asynchronous release<sup>9,12</sup><br>synaptic facilitation<sup>13</sup><br>vesicle replenishment<sup>14</sup><br>suppression of spontaneous release<sup>15</sup> |
| 3 | 3 | high<sup>2</sup> | yes<sup>3,4</sup> | 6.2 | synapses<sup>16</sup>, dendrites<sup>5,17</sup> | plasma membrane<sup>16</sup> | endocytosis<sup>17</sup> | postsynaptic GluR trafficking<sup>17</sup> |
| 3 | 6 | high | no<sup>3</sup>, yes<sup>4</sup> | 3.5 | synapses<sup>16</sup>, axons/dendrites<sup>5</sup> | plasma membrane<sup>16</sup> |  |  |
| 3 | 9**** | high<sup>2</sup> | yes<sup>3,4</sup> | 2.1 | axons<sup>5</sup> | plasma membrane |  |  |
| 3 | 10 | high<sup>2</sup> | no<sup>5</sup>, yes<sup>4</sup> | 0.9 | soma, axons<sup>5</sup> | plasma membrane<sup>5</sup> | exocytosis<sup>8</sup> | IGF-1 secretion<sup>8</sup> |
| 4 | 4 | none<sup>4,18-20</sup> | no<sup>3,4,20</sup> | 3.4 | axons/dendrites<sup>5</sup> |  | modulation<sup>4</sup> | inhibition of synaptotagmin-1-mediated fusion<sup>4</sup> |
| 5 | 5***** | intermediate<sup>1</sup>, low<sup>10</sup> | yes<sup>4</sup> | 9.5 | synapses, axons/dendrites<sup>5</sup> | synaptic vesicles | exocytosis<sup>10</sup> | synchronous release<sup>10</sup> |
| 6 | 8 | none<sup>4,5,21</sup> | no<sup>3,4</sup> | 0.0001 | not detected in brain<sup>5,22-24</sup><br>soma<sup>21</sup> | cytosol<sup>21</sup> | modulation<sup>4</sup> | inhibition of synaptotagmin-1-mediated fusion<sup>4</sup> |

\* *There are several divisions for the synaptotagmin family in literature. The one we adopt here is from Sudhof (2002).*
\*\* *Ca<sup>2+</sup> sensitivity.*
\*\*\* *Ca<sup>2+</sup>-dependent phospholipid binding.*
\*\*\*\* *The ratio of RNA reads per kilobase per million reads placed (RPKM) from brain vs. the maximal RPKM elsewhere in the body, calculated from quantitative transcriptome analysis (RNA-Seq) of 27 human tissue samples from 95 individuals (Fagerberg et al., 2014). Most synaptotagmins are more expressed in the brain (i.e. RPKM ratio >1) than elsewhere in the body, except for synaptotagmins 8 and 10, which have very low expression in the brain (RPKM <1).*
\*\*\*\*\* *Called synaptotagmin 5 in some studies (Li et al., 1995; Dean et al., 2012). Because of this confusion in the literature (Haberman et al., 2003), data from articles that did not make clear which convention was used (NCBI/Ensembl or naming based on papers from Thomas Sudhof's group) were not included in this table.*
\*\*\*\*\* *Called synaptotagmin 9 in some studies (Bhalla et al., 2005; Xu et al., 2007; Dean et al., 2012). Data from articles that did not make clear which convention was used were not included in this table.*
Refs.: <sup>1</sup>(Bhalla et al., 2005); <sup>2</sup>(Sugita et al., 2002); <sup>3</sup>(Li et al., 1995); <sup>4</sup>(Bhalla et al., 2008); <sup>5</sup>(Dean et al., 2012); <sup>6</sup>(Takamori et al., 2006); <sup>7</sup>(Geppert et al., 1994); <sup>8</sup>(Cao et al., 2013); <sup>9</sup>(Bacaj et al., 2013); <sup>10</sup>(Xu et al., 2007); <sup>11</sup>(Sugita et al., 2001); <sup>12</sup>(Luo et al., 2015); <sup>13</sup>(Jackman & Regehr, 2017); <sup>14</sup>(Liu et al., 2014); <sup>15</sup>(Luo & Sudhof, 2017); <sup>16</sup>(Butz et al., 1999).

Our analysis (**Figures 1–3**) and published data on genomic homology (Craxton, 2004, 2010) point to functional conservation of most synaptotagmin proteins in zebrafish. To address this question more rigorously, we looked at the Ca^2+^ binding pockets of the C2A and C2B domains. We focused specifically on whether zebrafish synaptotagmins could act as Ca^2+^ sensors by examining the amino acid residues responsible for Ca^2+^ coordination in the C2 domains.

### Conservation of crucial residues for Ca^2+^ binding

The interaction between synaptotagmins and Ca^2+^ is electrostatic in nature and creates an electrical switch for phospholipid binding (Ubach et al., 1998). In mammalian synaptotagmin 1, six negatively charged amino acid residues located in loops 1 and 3 of the C2A domain coordinate together the binding of three Ca^2+^ ions (**Figure 4A**), and five negative residues in loops 1 and 3 of the C2B domain coordinate the binding of two Ca^2+^ ions (**Figure 4B**). When not bound to Ca^2+^, the negative surface charge of these loops likely prevents close contact between the C2 domains and negatively charged phospholipid membranes. Binding of Ca^2+^ neutralizes these charges and promotes direct interaction between synaptotagmins and the cell membrane (or vesicular membrane, in the case of membrane-bound synaptotagmins, **Table 5**). Since the amino acid composition of these loops controls both the charge (Ubach et al., 1998) and exact shape (Dai et al., 2004; Qiu et al., 2017) of the Ca^2+^ binding pocket, it is therefore critical for the role of synaptotagmins as Ca^2+^ sensors for exocytosis.

**Figure 4.**
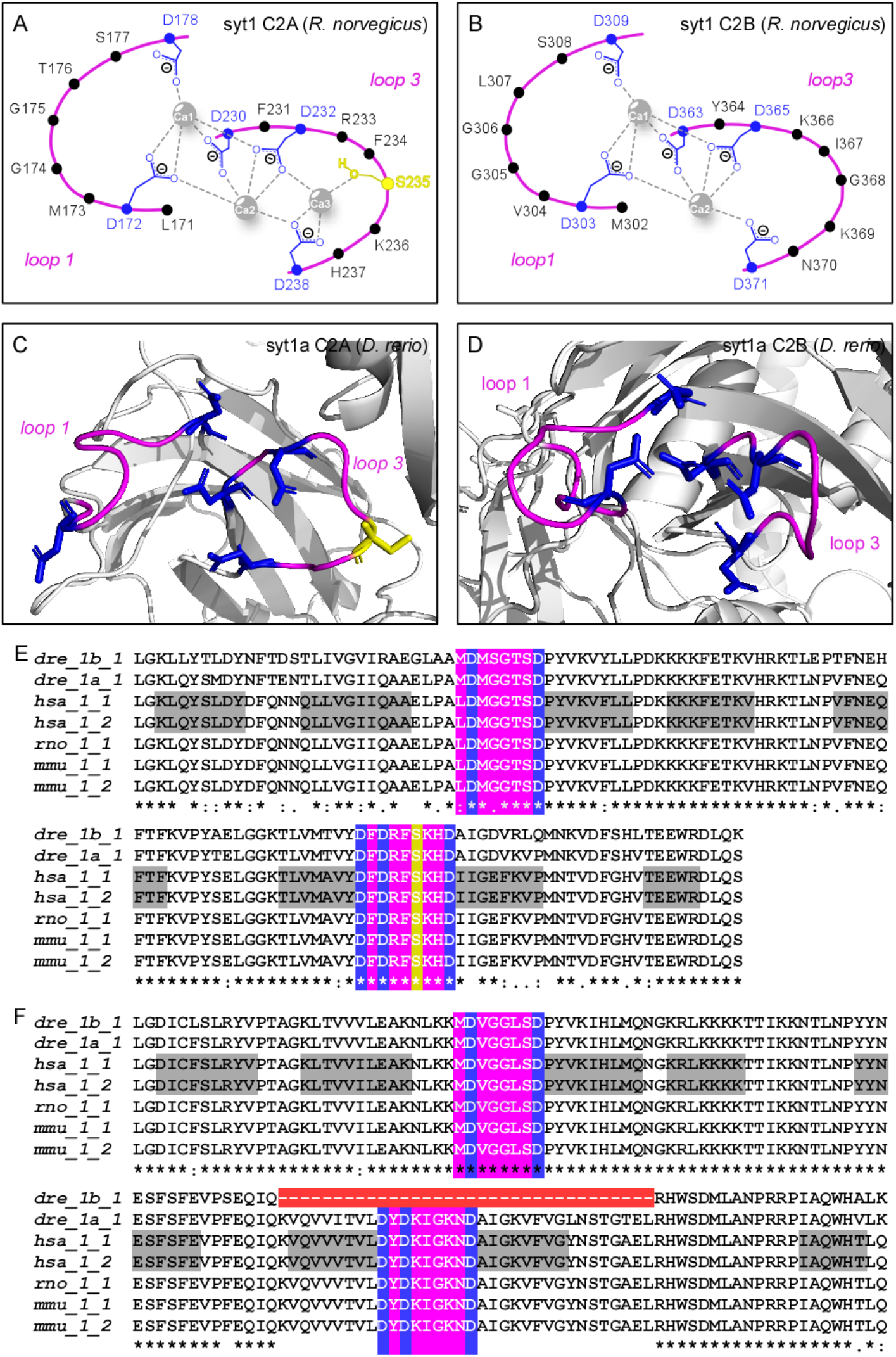
Homology of the functional domains of zebrafish synaptotagmin 1 paralogues. (A, B) Diagrams summarizing the three Ca^2+^-binding sites of the C2A domain (A) or the two Ca^2+^-binding sites of the C2B domain (B), coordinated by loops 1 and 3 (pink), of rat synaptotagmin 1. The negative residues responsible for the coordination of Ca^2+^ binding are shown in blue (for aspartate) or yellow (serine). Because the C2B domain lacks one negative residue (the serine in C2A), one of the Ca^2+^ binding sites is missing. Redrawn from Fernandez *et al*. (2001). (C, D) Ribbon models of the three-dimensional structures of the C2A (C) and C2B (D) domains of zebrafish synaptotagmin 1a generated with RaptorX. Loops 1 and 3 are shown in pink. Residues responsible for Ca^2+^ binding are shown in same colors as in (A,B). (E, F) Alignment of the amino acid sequences of the C2A (E) and C2B (F) domains of zebrafish synaptotagmin 1a and 1b (*dre_1a_1* and *dre_1b_1,* accession numbers in **Table 3**) with those of isoforms 1 and 2 of human (*hsa_1_1* and *hsa_1_2,* accession numbers in **Table 4**), rat (*rno_1_1:* NCBI# NP_001028852.2) and mouse (*mmu_1_1:* NCBI# NP_001239270.1 and *mmu_1_2:* NCBI# NP_001239271.1) synaptotagmin 1. Locations of the β-strands in the human sequence are highlighted in grey, after (Fuson et al., 2007); loops 1 and 3 are highlighted in pink, and negative residues responsible for Ca^2+^ binding are in blue (aspartates) or yellow (serine). Consensus between sequences is indicated below the alignments: “**\***”, identical residue; “:”, conservative replacement; “.”, semi-conservative substitution. All charged residues that coordinate Ca^2+^ in the C2A domain of mammalian synaptotagmins are conserved in both zebrafish paralogues, but 34 residues are missing in a region that includes two β sheets (β6 and β7) and loop 3 from the C2B domain of zebrafish synaptotagmin 1b (highlighted in red). This paralogue, therefore, is likely unable to bind Ca^2+^.

### C2 domains in Class 1 synaptotagmins

Ca^2+^ binding to the C2B domain of synaptotagmin 1 is crucial for synchronous release (Mackler & Reist, 2001; Mackler et al., 2002; Nishiki & Augustine, 2004; Shin et al., 2009; Bacaj et al., 2013; Lee et al., 2013), probably because the C2B domain has significantly higher sensitivity to Ca^2+^ (Bradberry et al., 2020) and phospholipid-binding activity (Bai et al., 2004; Li et al., 2006; van den Bogaart et al., 2012; Bradberry et al., 2020) than the C2A domain in this protein.

Indeed, all mutations in synaptotagmin 1 that cause diseases in humans target the C2B domain (Baker et al., 2018; Bradberry et al., 2020), while removal of the residues critical for Ca^2+^ binding in the C2A domain is relatively innocuous (Stevens & Sullivan, 2003).

Zebrafish synaptotagmin 1a has all 11 residues needed to coordinate Ca^2+^ binding at the same locations in both C2 domains as mammalian synaptotagmin 1, which render the 3-dimensional arrangement of the Ca^2+^ binding pockets very similar to that of its mammalian orthologues (**Figure 4C-D**). Alignment of the amino acid sequences of both C2 domains of zebrafish (dre), human (hsa), rat (rno) and mouse (mmu) orthologues confirms that these regions are highly conserved in synaptotagmin 1a (**Figure 4E-F**). This paralogue is therefore likely to function as a Ca^2+^ sensor. On the other hand, synaptotagmin 1b lacks a stretch of 34 amino acids around loop 3 of the C2B domain (area highlighted in red in the alignment in **Figure 4F**). This truncation probably renders this paralogue unable to coordinate Ca^2+^ binding and to function as a sensor for exocytosis.

**Table 6** summarizes the homology of the Ca^2+^ binding motifs in both C2 domains between zebrafish, human, rat and mouse orthologues. Interestingly, the Ca^2+^ binding motifs of synaptotagmins 2a and 2b are more homologous to mammalian synaptotagmin 1 than to synaptotagmin 2. The C2B domains of the zebrafish proteins are identical to those of mammalian synaptotagmin 1, while mammalian synaptotagmin 2 has a conservative aspartate (D)-to-glutamate (E) substitution in the C2B domain. Such a substitution is also present in mammalian synaptotagmin 3 (and in all class 3 orthologues, **Table 6**) and may lead to collapse of the last Ca^2+^ binding site of this domain (Sutton et al., 1999). Therefore, zebrafish synaptotagmins 2a and 2b could probably bind the same number of Ca^2+^ ions as synaptotagmins 1 and 1a, which is one Ca^2+^ ion more than mammalian synaptotagmin 2. Also, since the C2B domain is more important for synaptotagmin 1 function in synchronous release (Mackler et al., 2002; Nishiki & Augustine, 2004), it is likely that synaptotagmin 2a and 2b have similar properties as synaptotagmin 1. Notably, synaptotagmin 2b underlies synchronous release in the zebrafish neuromuscular junction (Wen et al., 2010).

**Table 6.**
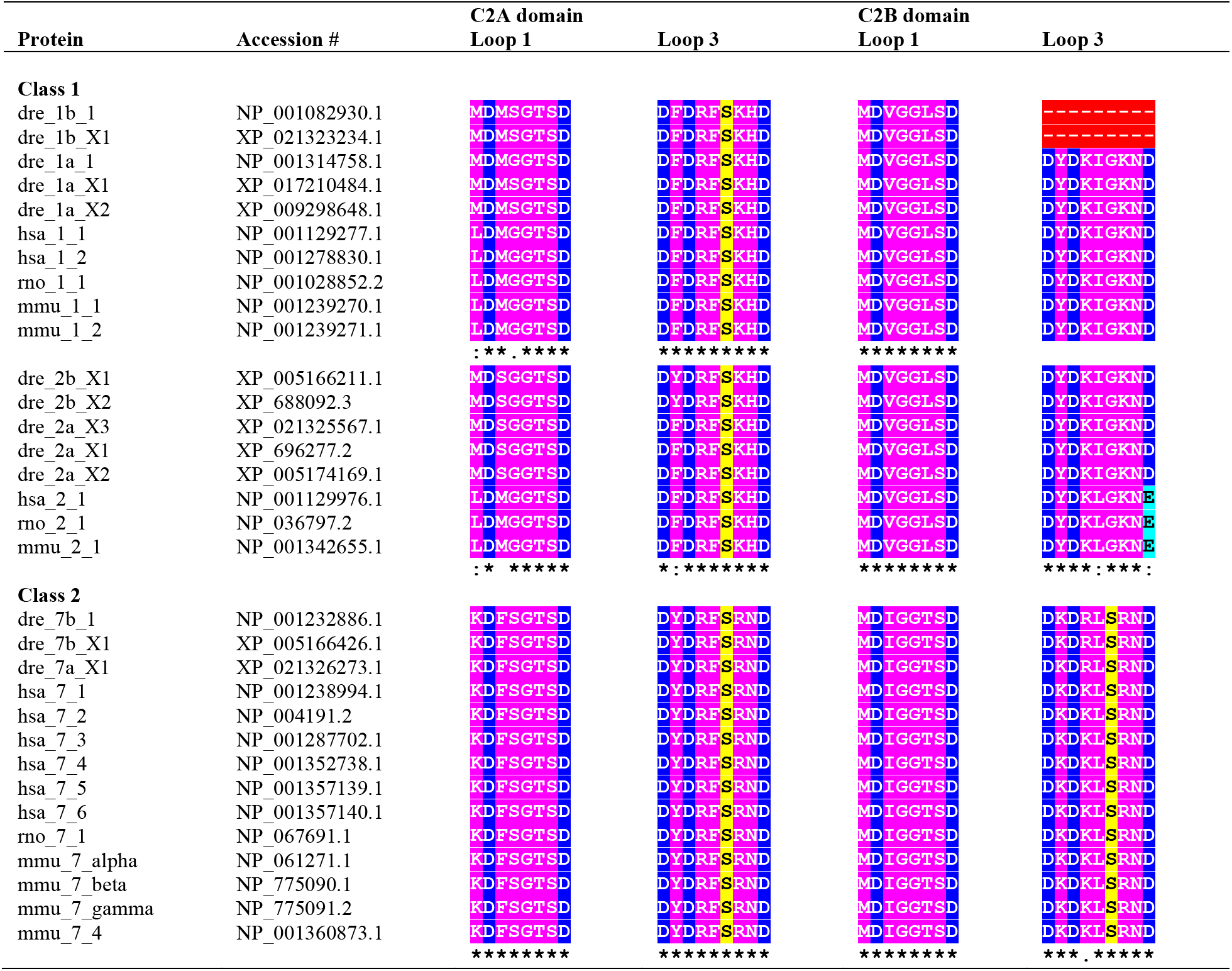
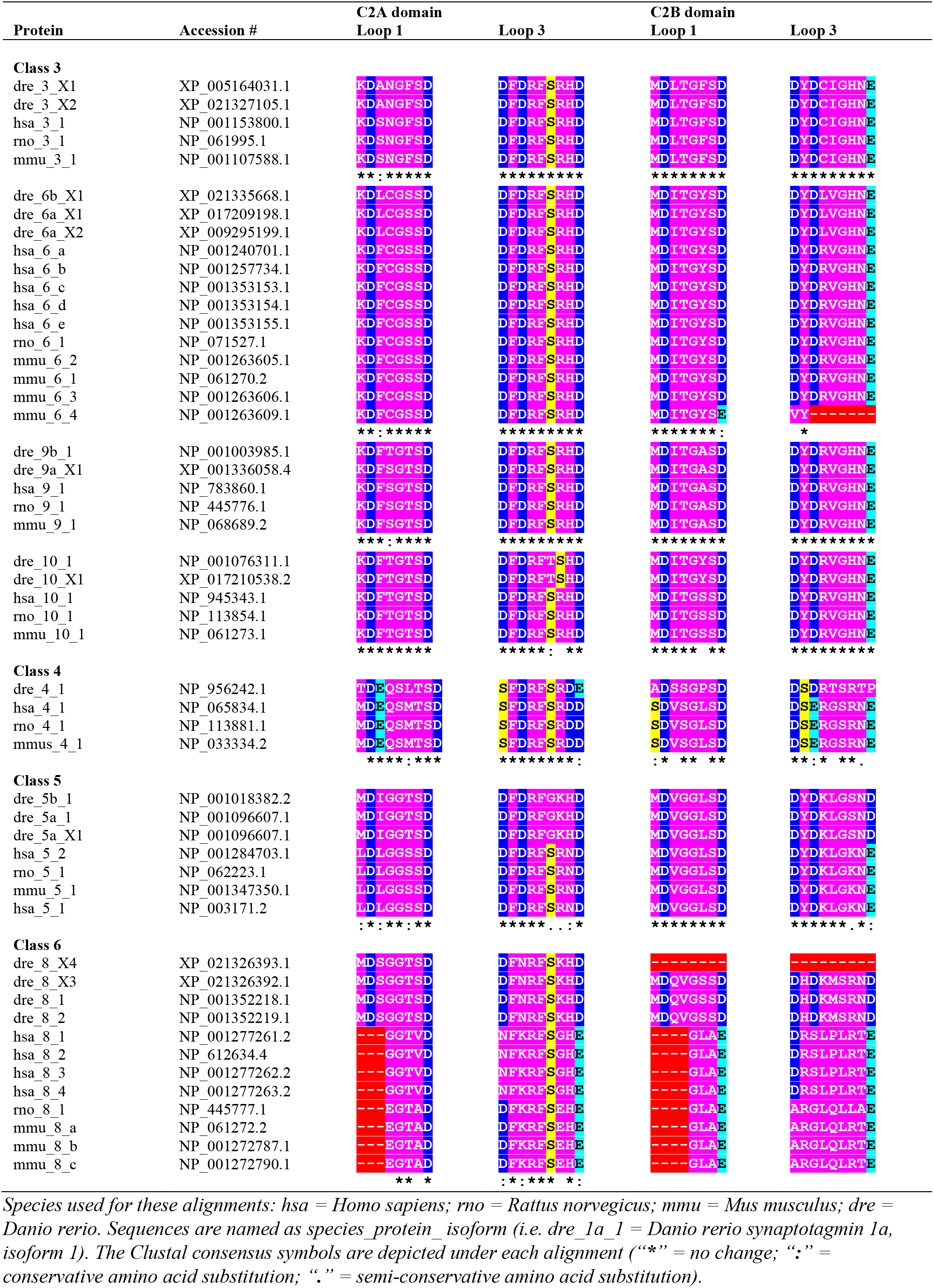
Homology of the Ca^2+^ binding motifs.

### C2 domains in Class 2 and 3 synaptotagmins

Zebrafish class 2 synaptotagmins have only one semi-conservative amino acid substitution at a non-essential location in the C2B domain in relation to their mammalian orthologues. There are no substitutions in the C2A domain, which has been deemed more important than the C2B domain for controlling exocytosis in this class (Xue et al., 2010; Bacaj et al., 2013; Voleti et al., 2017). This extreme degree of conservation is highly suggestive of functional preservation (**Table 5**). In fact, the 3-dimensional model of the C2 domains of synaptotagmin 7b aligns well with the crystal structure of the C2 domains of rat synaptotagmin 7 (PDB: 6ANK, Voleti et al., 2017)(**Figure 5A**). It is interesting to note that the C2B domain of mammalian synaptotagmin 7 can bind 3 Ca^2+^ ions instead of 2, due to the presence of a serine residue in loop 3 (Xue et al., 2010). This makes the C2B domain of these proteins highly similar to the C2A domain as far as Ca^2+^ coordination is concerned, although its functional significance is yet unknown (Xue et al., 2010). This negative residue is also present in the C2B domain of zebrafish class 2 synaptotagmins (highlighted in yellow in **Table 6**).

**Figure 5.**
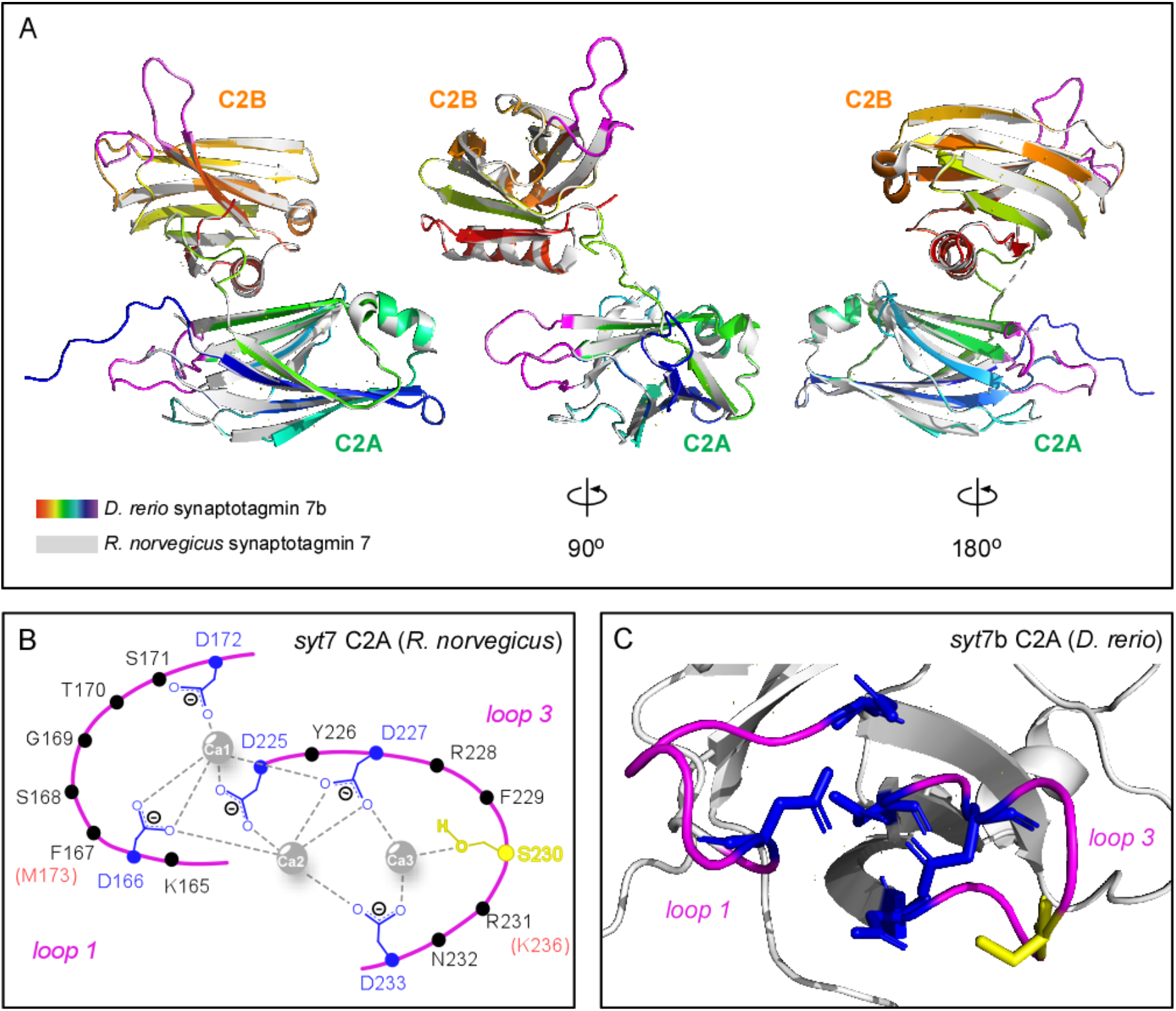
Homology of the functional domains of zebrafish synaptotagmin 7b. (A) Ribbon model of the 3-dimensional structure of zebrafish synaptotagmin 7b, isoform 1 (rainbow colors) generated with RaptorX from the amino acid sequence (isoform 1, accession number in **Table 3**) and superposed on the crystal structure of the C2 domains of rat synaptotagmin 7 (shown in grey, PDB: 6ANK, Voleti et al., 2017). Figure is rotated 90 (center) and 180 degrees (right) to display the loops responsible for Ca^2+^ binding in the C2 domains (loops 1 and 3) of the rat synaptotagmin (pink). (B) Diagram summarizing the three Ca^2+^-binding sites of the C2A domain, coordinated by loops 1 and 3 (pink), of rat synaptotagmin 7. The negative residues responsible for the coordination of Ca^2+^ binding are shown in blue (for aspartate) or yellow (serine). Redrawn from Voleti *et al*. (Fernandez et al., 2001; 2017). (C) Ribbon model of the three-dimensional structure of the C2A domain of zebrafish synaptotagmin 7b generated with RaptorX. Loops 1 and 3 are shown in pink. Residues responsible for Ca^2+^ binding are shown in same colors as in (B).

Since synaptotagmin 7 was implicated in Ca^2+^-dependent asynchronous release in mammals (Bacaj et al., 2013; Ablain et al., 2015), probably due to the higher affinity of its Ca^2+^-bound C2A domain for phospholipids (Voleti et al., 2017), it is likely to perform the same role in the zebrafish. Indeed, synaptotagmin 7b was reported to control asynchronous release at the zebrafish neuromuscular junction (Wen et al., 2010). Other roles for mammalian synaptotagmin 7 in synaptic transmission are listed in **Table 5**.

Both class 2 and class 3 synaptotagmins show two crucial amino acid substitutions in their C2A domains in relation to class 1 synaptotagmins that, in mammals, determine their much higher affinity to phospholipid membranes: the methionine (M)-to-phenylalanine (F) substitution at the third position of loop 1 and the lysine (K)-to-arginine (R) substitution at the seventh position of loop 3 (Voleti et al., 2017). Although these are conservative replacements, they seem to highly impact Ca^2+^-dependent phospholipid binding and show cooperative effects: the highest affinity is found when both substitutions are present (i.e. synaptotagmins 7, 7a, 7b, 6, 9a, 9b and mammalian 10). Zebrafish synaptotagmin 10 is an exception here: it has as negatively charged residue (serine) at the seventh position of loop 3 and no serine at the sixth position, which may impact its ability to coordinate Ca^2+^ binding.

### C2 domains in Class 4 and 6 synaptotagmins

Class 4 and 6 synaptotagmins are the most divergent (**Figure 3**, **Table 6**). These proteins are unable to bind Ca^2+^ in mammals and function therefore as inhibitory synaptotagmins in these species (von Poser et al., 1997; Dai et al., 2004; Hui et al., 2005). Zebrafish synaptotagmin 4 misses more aspartates than the mammalian orthologues in the C2A domain and may not bind Ca^2+^ there, because the two missing aspartates coordinate together all Ca^2+^ binding sites in this domain. Since there is evidence that all three Ca^2+^ ions are required for binding with syntaxin (Ubach et al., 1998), this could render this synaptotagmin unable to drive exocytosis. Of note, the amino acid substitutions in the C2A domain are semi-conservative (aspartate (D)-to-serine (S)) or conservative (aspartate (D)-to-glutamate (E)), so there is a chance that these negatively charged residues could still create Ca^2+^ binding sites ((Dai et al., 2004), but see (von Poser et al., 1997; Wang & Zhang, 2017)). The C2B domain of zebrafish synaptotagmin 4 has more aspartates than its mammalian orthologues and may still be able to bind at least one Ca^2+^ ion according to the model in **Figure 4A-B**(Fernandez et al., 2001).

Whether a synaptotagmin will bind Ca^2+^ efficiently depends on both the amino acid sequence and on the 3-dimensional structure of loops 1 and 3 in each C2 domain (i.e. the orientation of the negative residues in space) (Dai et al., 2004). This 3-dimensional structure, in turn, is determined by the amino acid composition of the loops themselves and of the flanking regions that interact electrically with the loops (Qiu et al., 2017). For instance, although rat and *Drosophila* synaptotagmin 4 have exactly the aspartate (D)-to-serine (S)) substitution in loop 1 of the C2A domain, only in the latter does it function as a Ca^2+^ sensor, presumably because loop 1 in rat has a different 3-dimensional structure (Dai et al., 2004). This is an example of homology without orthology; clearly, this protein suffered a change in function during evolution. It remains to be determined whether zebrafish synaptotagmin 4 can still function as a Ca^2+^ sensor.

While mammalian synaptotagmin 8 misses significative portions of loop 1 in both C2A and C2B domains, at least three isoforms of zebrafish synaptotagmin are not only intact but have most Ca^2+^ coordinating residues. The only exception is isoform X4, which is truncated. The first aspartate residue missing in loop 3 of the C2A domain in all orthologues coordinates binding of all 3 Ca^2+^ ions in synaptotagmin 1 (**Figure 4**); a study with mammalian synaptotagmins 1 and 2 suggests that its absence may lead to collapse of the whole Ca^2+^ binding pocket in this domain (Stevens & Sullivan, 2003). It is therefore unlikely that the C2A domain of zebrafish synaptotagmin 8 can bind Ca^2+^, despite having more conserved residues in relation to other synaptotagmins than its mammalian orthologues. The C2B domain, however, has all 5 aspartates in this species, so it may still potentially coordinate Ca^2+^ binding. Further studies will be needed to address this issue.

### C2 domains in Class 5 synaptotagmins

As discussed earlier, zebrafish synaptotagmin 5a and 5b fall between mammalian class 1 and 5. We kept them in class 5 for this analysis to keep the nomenclature consistent. The low overall genomic homology between zebrafish synaptotagmin 5a and 5b is also reflected in the amino acid composition of their Ca^2+^ coordinating loops. Both synaptotagmin 5a and 5b lack the serine necessary to close the C2A binding pocket and coordinate Ca^2+^ binding in the third site of this domain (Qiu et al., 2017). However, the same semi-conservative serine (S)-to-glycine (G) substitution in human synaptotagmin 5 does not compromise its three Ca^2+^ binding sites. Rather, this substitution leads only to a slight conformational change in loop 3 of the C2A domain, which renders the Ca^2+^ binding pocket more open and decreases Ca^2+^ affinity, making synaptotagmin 5 more synaptotagmin 1-like (Qiu et al., 2017). Further, the zebrafish synaptotagmins have all aspartates in the C2B domain, whereas the mammalian orthologues have a conservative aspartate (D)-to-glutamate (E) substitution in relation to other isoforms, similar to the one found in class 3 synaptotagmins, and may actually bind less Ca^2+^ ions there (Sutton et al., 1999). Therefore, the Ca^2+^ binding motifs of zebrafish synaptotagmins 5a and 5b may be more homologous to those of class 1 synaptotagmins than to those of other class 5 synaptotagmins. These results, added to the fact that mammalian synaptotagmins 1, 2 and 5 were shown to be involved in synchronous release in central synapses (Xu et al., 2007), make zebrafish 5a and 5b good candidates for mediating fast exocytosis.

In summary, the results presented thus far point to a generally good functional homology between zebrafish and mammalian synaptotagmins at the protein level, especially for classes 1, 2, 3 and 5. We next concentrated on the zebrafish isoforms that could likely serve as Ca^2+^ sensors for neurotransmitter release in the retina.

### Expression and distribution of synaptotagmin candidates for Ca^2+^-dependent exocytosis in the zebrafish retina

Since class 3 synaptotagmins in mammals are mostly involved in processes other than vesicle release proper, such as endocytosis and non-synaptic secretion (**Table 5**), we set out to determine the retinal expression of *syt1, syt2, syt5* and *syt7* paralogues. RT-PCR cDNA fragments of all these genes were found in the zebrafish brain, eye, and retina, except for *syt1b* (**Figure 6** and **Table 7**). This paralogue was not detected in the eye or retina but was present in brain.

**Table 7.** Expression of *syt* mRNAs in different tissues assayed by RT-PCR.

| Zebrafish gene | Brain | Eye | Retina |
| --- | --- | --- | --- |
| *syt1a* | + | + | + |
| *syt1b* | + | - | - |
| *syt2a* | + | + | + |
| *syt2b* | + | + | + |
| *syt5a* | + | + | + |
| *syt5b* | + | + | + |
| *syt7a* | + | + | + |
| *syt7a<sub>splice</sub>* | + | + | + |
| *syt7b* | + | + | + |

**Figure 6.**
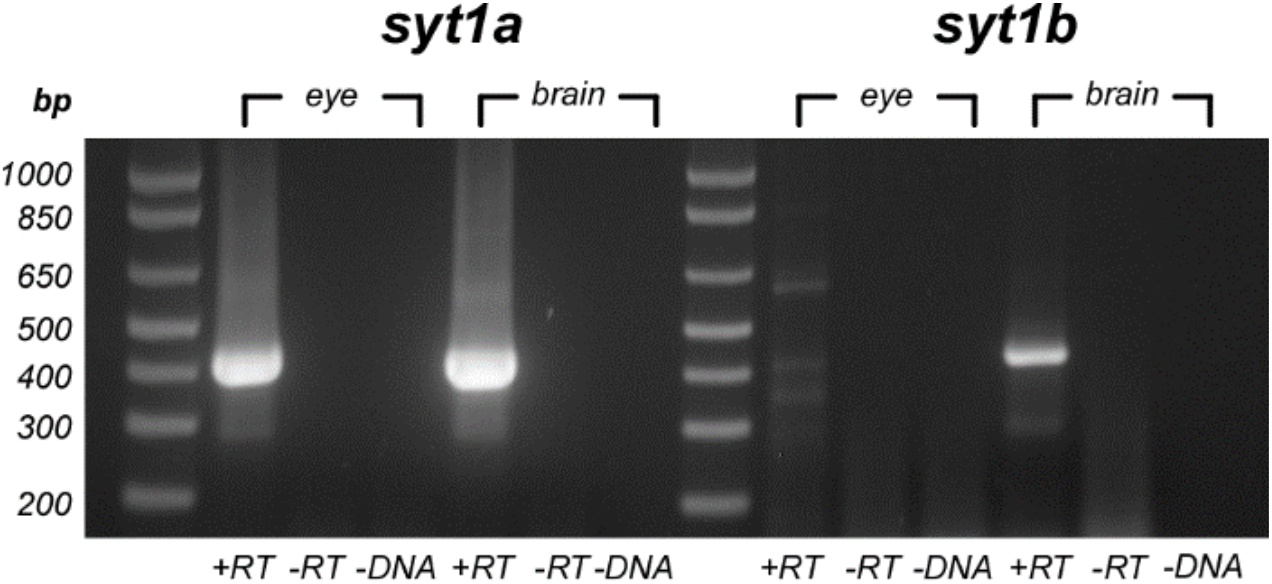
Retinal expression and distribution of zebrafish *syt1* paralogue genes. Agarose gel electrophoresis of RT-PCR cDNA products amplified from total RNA from adult zebrafish brain or eye using primers for *syt1a* and *syt1b* paralogues (**Table 1**). The sizes of the molecular markers are indicated on the left (*bp*: base pairs). *Syt1a* is present in both brain and eye, whereas *syt1b* is only present in the brain. The different lanes for each tissue represent tissue RNA + reverse transcriptase (+*RT*), plus two controls: tissue RNA without reverse transcriptase (-*RT*), and tissue RNA with no DNA (-DNA).

We also identified expression of a new splice variant of *syt7a* in zebrafish eye (named “*syt7a_splice*” in **Table 7**). This variant has close homology with *syt7a*-like sequences in bony fishes (ex: XP_016324438.1) but misses 774 base pairs in relation to variant XM_021470598.1, which codes for a long version of the protein (653 amino acids, **Table 3**). Synaptotagmin 7a_splice should therefore be around 395 amino acids long. In mammals, the L1 region (**Figure 2**) of synaptotagmin 7 undergoes extensive alternative splicing (Sugita et al., 2001). This leads to dramatic changes in protein structure and several protein isoforms of different lengths (**Table 4**), but this truncation at a linker region is of minor consequence for protein function. The alternatively spliced sequence also codes for L1 and should therefore give rise to a fully functional protein. The sequenced fragment for *syt7a_splice* is in the *Appendix*.

To study protein distribution in the adult zebrafish retina in more detail, we assayed mRNA expression for the different isoforms by *in situ* hybridization on retinal slices. Riboprobes were designed using ATG as the 5’ start if sequences were divergent in this area. Alternate primer locations were used if isotypes were not divergent. Since we identified a splice variant in *syt7*a, its riboprobe was located in the 3’ UTR to ensure that expression of all *syt7*a variants were detected.

We tested two isotype-specific riboprobes for synaptotagmin 1a (**Table 2**): a shorter probe that targeted only the coding sequence (CDS) and a longer riboprobe that included the 3’-UTR and could presumably enhance signal in case of low expression levels, as well as further validate the expression pattern seen with original probe (**Figure 7A-B**). Both yielded similar expression patterns, albeit the signal-to-noise ratio was better with the shorter CDS probe. *Syt1a* mRNA could be detected in three locations: (i) the outer plexiform layer (*OPL*), where photoreceptors contact bipolar cells and horizontal cells; (ii) the inner border of the inner nuclear layer (*INL*), where the somas of cone-driven bipolar cells and amacrine cells are located; and (iii) around somas in the ganglion cell layer (*GCL*).

**Figure 7.**
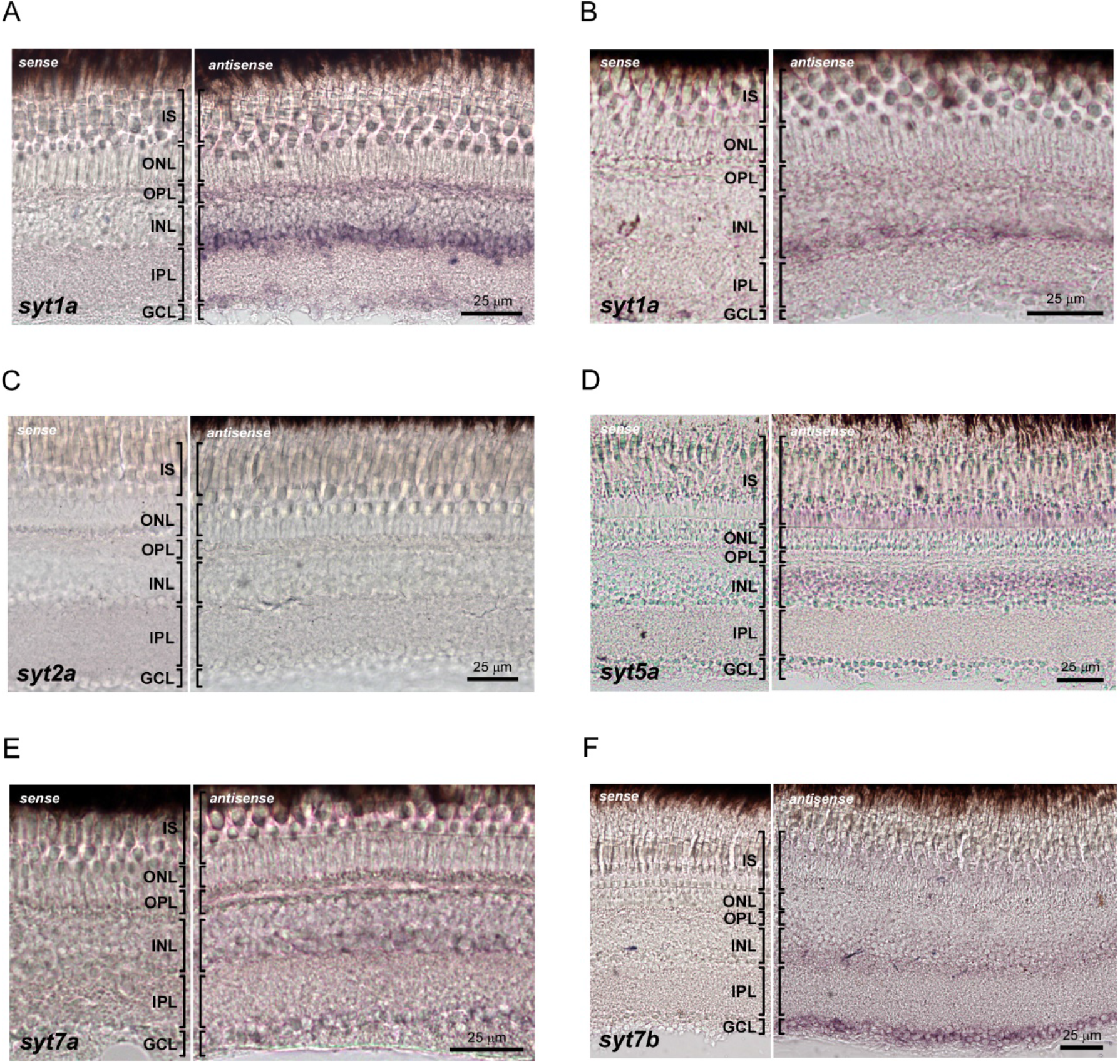
Retinal expression and distribution of zebrafish *syt* mRNAs assayed by *in situ* hybridization. For all panels, the left panel shows the negative control with a sense probe. *IS*: photoreceptor inner segments; *ONL*: outer nuclear layer; *OPL*: outer plexiform layer; *INL*: inner nuclear layer; *IPL*: inner plexiform layer; *GCL*: ganglion cell layer. (A, B) Detection of *syt1a* mRNA using a short CDS riboprobe (A) and a longer 3’-UTR riboprobe (B). *Syt1a* is expressed at highest levels by cells in the *OPL*, inner border of the *INL* and in the *GCL*. (C) *Syt*2a mRNA was undetectable by a short CDS type-specific riboprobe. An additional riboprobe (perhaps larger fragment in the UTR specific to *syt*2a) needs to be tested. (D) S*yt5a* mRNA is found at highest levels by inner segments of small cones located at the border with the *ONL*, and by cells at the outer half of the *INL*. (E) *Syt7a* mRNA can be detected at the *OPL*, inner half of the *INL* and *GCL*. (F) S*yt7b* is expressed at highest levels by cells at the inner border of the *INL* and by cells in the *GCL*.

Although we sequenced a fragment consistent with *syt2a* in the zebrafish retina by PCR, our short CDS riboprobe did not yield any signal (**Figure 7C**). This result could mean that either the *syt2a* paralogue is not transcribed, the products of this gene could be targeted for nonsense mediated decay, and/or that the probe was too short to hybridize strongly. Further experiments with a longer isotype-specific probe are needed to disambiguate this result. It may be necessary to confirm that the full-length *syt*2a gene is indeed found in zebrafish retinal tissue. If positive, specific probes, located in either the 3’ or 5’ UTR should be generated.

Weak labeling for *syt5a* mRNA was seen in the inner segments (IS) of short cones and in the outer half of the INL, were the cell bodies of horizontal cells and of mixed-input bipolar cells (i.e. bipolar cells that contact both rods and cones) and cone-driven bipolar cells are located (**Figure 7D**). The expression pattern of *syt7a* was like that of *syt1a*, which could mean that these synaptotagmins 1a and 7a are expressed in the same neurons in the zebrafish (**Figure 7E**). Finally, *syt7b* was detected at the inner border of the INL and in the GCL (**Figure 7F**). The differential expression of *syt1a* and *syt5a* suggest that these isoforms may control synchronous release in distinct neurons in the zebrafish retina, while *syt7a* and *syt7b* would mediate asynchronous release, vesicle replenishment or other Ca^2+^ dependent processes, probably in different neuronal populations as well.

## DISCUSSION

In this study, we took advantage of bioinformatics to address the similarity between zebrafish and human synaptotagmins. We focused on synaptotagmin 1-10 as their roles are better defined and synaptotagmins 11-17 are unlikely candidates to mediate Ca^2+^ exocytosis in the central nervous system (von Poser et al., 1997; Fukuda, 2003a, b; Bhalla et al., 2008; Craxton, 2010). Because of whole genome duplication, zebrafish generally have two paralogues for each human orthologue, except for *syt*3, *syt*4, *syt*8 and *syt*10 (Craxton, 2004, 2010). Despite duplication and the evolutionary distance, we find that most zebrafish synaptotagmins share common features with mouse, rat and human orthologues, including key motifs involved in Ca^2+^ binding.

Accordingly, we find that *syt*1a, *syt*2a, *syt*2b, *syt*5a, *syt*5b, *syt*7a and *syt*7b genes are expressed in the zebrafish brain, eye and retina. From these, we successfully detected mRNA for *syt*1a, *syt*5a, *syt*7a and *syt*7b in retinal tissue with distinct distribution patterns. This differential expression suggests that these synaptotagmins may control various aspects of the vesicle cycle in distinct subsets of neurons. Overall, these results highlight that zebrafish will be an outstanding model to study the role of synaptotagmins in synaptic transmission in the retina.

### Homology of zebrafish synaptotagmins

In the present study, we characterized genetic and protein similarities between zebrafish and human synaptotagmins (**Figures 1–3**). Importantly, zebrafish synaptotagmins cluster mostly into the same classes as their mammalian counterparts (**Figure 3**), except synaptotagmins 5a and 5b, which may be more homologous to class 1 synaptotagmins. In addition, zebrafish synaptotagmins show strong similarities to their mammalian counterparts in terms of their Ca^2+^ binding motifs (**Figures 4**-**5**, and **Table 6**). Notable exceptions are zebrafish synaptotagmin 1b, which lacks 34 amino acid residues in its C2B domain, and synaptotagmin 8, which has more crucial residues for Ca^2+^ binding in both C2 domains than the mammalian synaptotagmin 8.

Synaptotagmin 1b is unlikely to act as a Ca^2+^ sensor for exocytosis. In mammals (Dai et al., 2004; Bhalla et al., 2008; Xue et al., 2010; Voleti et al., 2017), Ca^2+^ binding to the C2B domain of synaptotagmin 1 is crucial for synchronous release (Mackler & Reist, 2001; Mackler et al., 2002; Nishiki & Augustine, 2004; Shin et al., 2009; Bacaj et al., 2013; Lee et al., 2013), probably because the C2B domain has significantly higher sensitivity to Ca^2+^ (Bradberry et al., 2020) and phospholipid-binding activity (Bai et al., 2004; Li et al., 2006; van den Bogaart et al., 2012; Bradberry et al., 2020) than the C2A domain in this protein. Indeed, all mutations in synaptotagmin 1 that cause diseases in humans target the C2B domain (Baker et al., 2018; Bradberry et al., 2020), while removal of the residues critical for Ca^2+^ binding in the C2A domain is relatively innocuous (Stevens & Sullivan, 2003).

One could wonder why the *syt1b* gene is expressed at all, since its product lacks the ability to effectively bind Ca^2+^. Synaptotagmins can bind other synaptotagmins and t-SNARES in a Ca^2+^-independent manner, thereby displacing the Ca^2+^-sensitive isoforms and impeding direct contact between them and their targets (Bhalla et al., 2008). It is possible that synaptotagmin 1b may be one such inhibitory protein that regulates the activity of other synaptotagmins in the zebrafish brain, like mammalian synaptotagmins 4 and 8 (von Poser et al., 1997; Dai et al., 2004; Hui et al., 2005; Bhalla et al., 2008).

That said, it is important to note that sequence analysis alone predicts Ca^2+^ binding properties rather poorly. For example, despite the high homology of their C2 Ca^2+^ binding pockets, drosophila synaptotagmin 4 is Ca^2+^ sensitive, while its mammalian orthologue is not (Dai et al., 2004; Bhalla et al., 2008). Given that the differences in amino acid composition of the C2 domains of Ca^2+^-sensitive synaptotagmins are rather subtle, the basis for their Ca^2+^ sensitivity and functional differences remain poorly understood (Xue et al., 2010; Voleti et al., 2017). For both zebrafish synaptotagmin 1b and 8 further experiments are thus necessary to determine whether they may or may not function as Ca^2+^ sensors.

### Synaptotagmins expressed in the zebrafish retina

Although zebrafish is a powerful model organism, the duplication of its genome can often complicate the identification of homologous genes and their isotypes. Frequently, the available sequence data is not mapped, completely validated, or correctly named in the various databases. This bioinformatic study allowed us to examine the current genomic and mRNA sequences and ensure that current nomenclature for these various synaptotagmins are accurately based on their mammalian orthologues. With focus on those synaptotagmins that are likely candidates for mediating Ca^2+^-dependent exocytosis in the retina, we were thus able to confidently identify appropriate sequence data for *syt*1a/b, *syt*2a/b, *syt*5a/b and *syt*7a/b. These sequences were then used to design RT-PCR primers and ultimately riboprobes for identification of expressed synaptotagmins in the retina.

We find using RT-PCR that 7 of the 8 studied paralogues are present in retina, the exception being *syt1*b. This raises the possibility that the various synaptotagmins might make differential contributions to retinal physiology. Of these, we could only detect mRNA for *syt*1a, *syt*5a, *syt*7a and *syt*7b by *in situ* hybridization. This divergence might reflect that very short CDS riboprobes needed to guarantee isotype-specificity may limit detection. Longer riboprobes containing sequences from UTRs, which will maintain specificity and hopefully improve signal-to-noise ratio, will be needed to fully address this issue. Alternatively, it could be that some of the fragments amplified by RT-PCR (i.e from *syt*2a and *syt*2b) do not reflect the presence of a full-length gene in the retina. In this case, zebrafish and mammalian retinas could use different sets of synaptotagmins to control vesicle release in ribbon-type synapses of the OPL, since synaptotagmin 2 was reported in the rat and mouse OPL (Ullrich & Sudhof, 1994; Fox & Sanes, 2007), albeit in the mouse it seems to be excluded from photoreceptor terminals (Fox & Sanes, 2007). Future experiments will have to address this.

The expression of *syt*1a in the OPL is consistent with the immunofluorescence data in rat (Ullrich & Sudhof, 1994) and with the finding that synaptic release from photoreceptors in the mouse is largely controlled by synaptotagmin 1 (Grassmeyer et al., 2019). Additionally, the strong *syt*1a signal detected in the inner half of the INL and in the GCL is very similar to the one for *syt*1 mRNA in the mouse (Fox & Sanes, 2007). Together, these results suggest that synaptotagmin 1a may have analogous functions in the zebrafish and control synchronous vesicle release both in ribbon synapses of photoreceptors and in conventional synapses of the retina, such as those of amacrine cells and ganglion cells.

While synaptotagmin 7 was shown to be involved in types of asynchronous release both at conventional (Bacaj et al., 2013; Luo & Sudhof, 2017) and ribbon-type synapses of retinal bipolar cells (Luo et al., 2015), the localization of mRNA signal for *syt*7a and *syt*7b in our study is not consistent with this role in zebrafish bipolar cells. Rather, it suggests that these paralogues are expressed mostly in amacrine cells (inner half of the INL) and ganglion cells (GCL). The weak *syt*7a signal we find in the zebrafish OPL suggests that synaptotagmin 7a may be additionally be present in photoreceptors. However, the role(s) of these proteins in the zebrafish retina still need elucidating, since synaptotagmin 7 was associated with other aspects of neurotransmission, such as vesicle replenishment (Liu et al., 2014) and synaptic facilitation (Jackman et al., 2016; Jackman & Regehr, 2017).

Finally, the expression pattern of *syt*5a is consistent with cone photoreceptors (ONL and inner segments of short cones) and horizontal and bipolar cells (outer half of the INL). To our knowledge, this pattern does not resemble any data from mammals. This, added to the similarities between the products of zebrafish *syt*5 paralogues with both class 1 and class 5 synaptotagmins in mammals warrants further investigation regarding whether these synaptotagmins are directly involved in neurotransmission.

## Acknowledgments

This work was supported by a NIH RO1 grant from NEI (EY003821 to GGM/LPW). We thank Miaomiao He for assistance with protein structure modeling and David Zenisek and Howard Sirotkin for helpful discussions and/or comments on the manuscript.

## APPENDIX

## 1. cDNA fragment of *syt7a_splice (264 nt)*

*ISH primer sequences are shown in bold font*.

**ATGTATCTCAACAGGGAGGAGGAGTA**CAGCAAAGGCTCCATCTCTTTGAGCGTGC TGCTGGTGTCGTTGGCGGTAACTGTGTGTGGGGTTTGGCTGGTGGCTCTCTGTGGCGTCTGTGGATGGTGTCAACGCAAGCTGGGGAAGAGGAATAAACCCGGAGTGGAGTCCGTCGGGTCTCCAGATTCAGGAAGAGGAAGAGGGGAGAAAAAAGCCATCAATAGGA ATATGGGGAATAAGCCAG**CCAACAGTCCTAAGGGTCAGCCG**

## 2. BLASTN alignment to *Sinocyclocheilus anshuiensis* synaptotagmin-7-like (LOC107674837), transcript variant X1 (95% identity)

*Zebrafish syt7a_splice sequence is shown as “query”*.

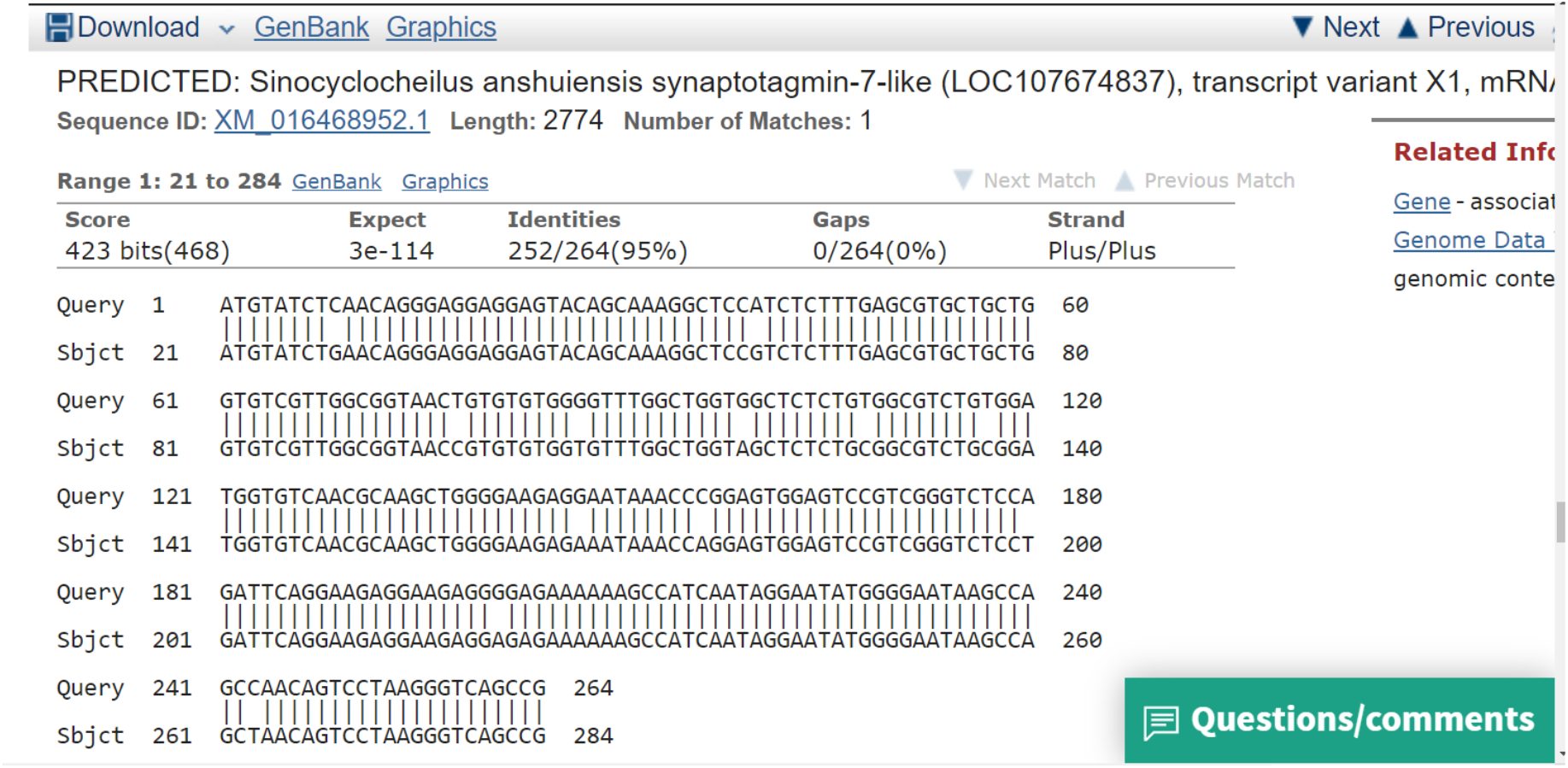

## 3. Genomic location of amplified sequence (not to scale)

Boxes represent exons of the *syt*7a gene in the long and splice variants.

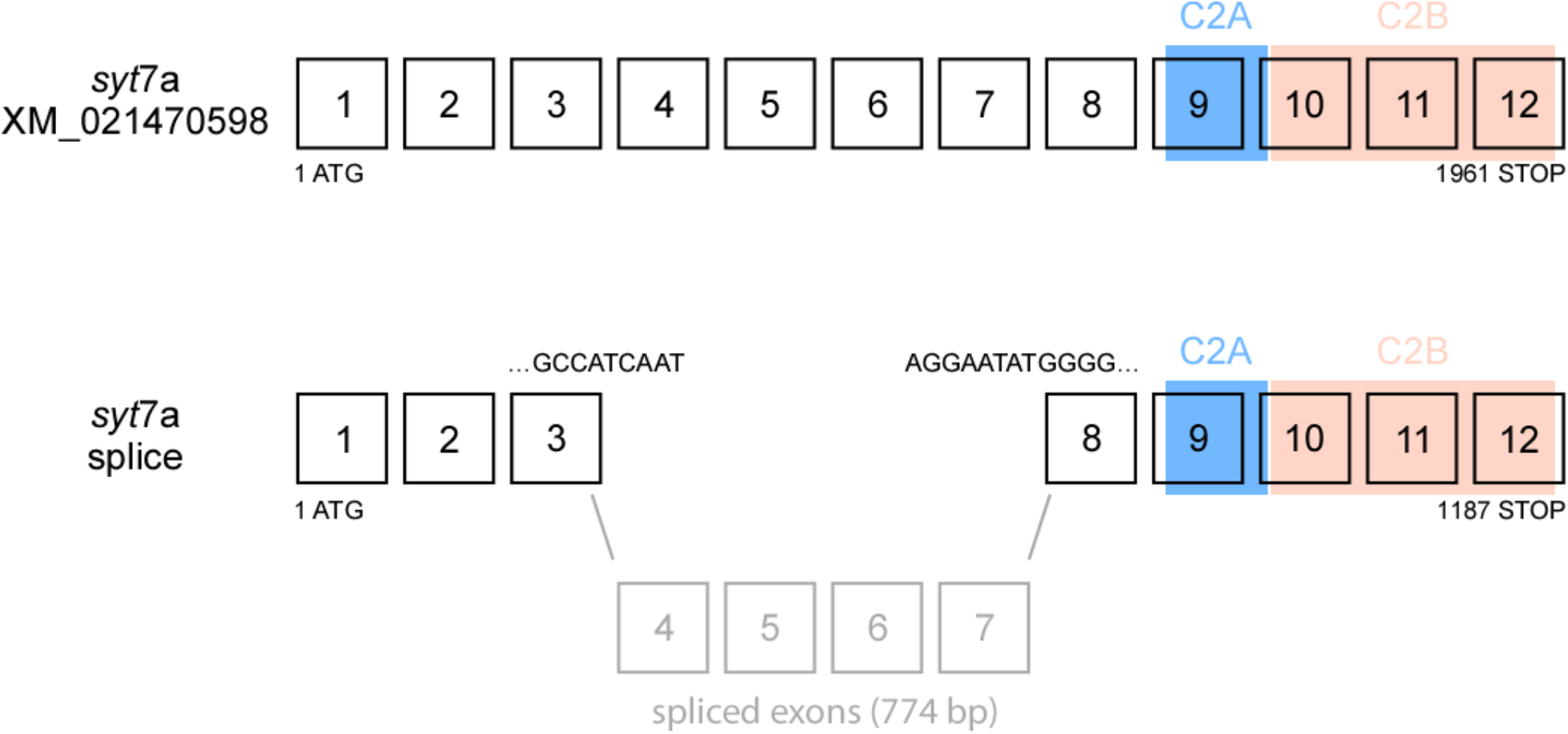

## 4. BLASTx alignment of the product of *syt* 7a_splice to *Sinocyclocheilus anshuiensis* synaptotagmin-7-like (XP_016324438.1), isoform X1 (98% identity)

*Zebrafish syt7a_splice sequence is shown as “query”.*

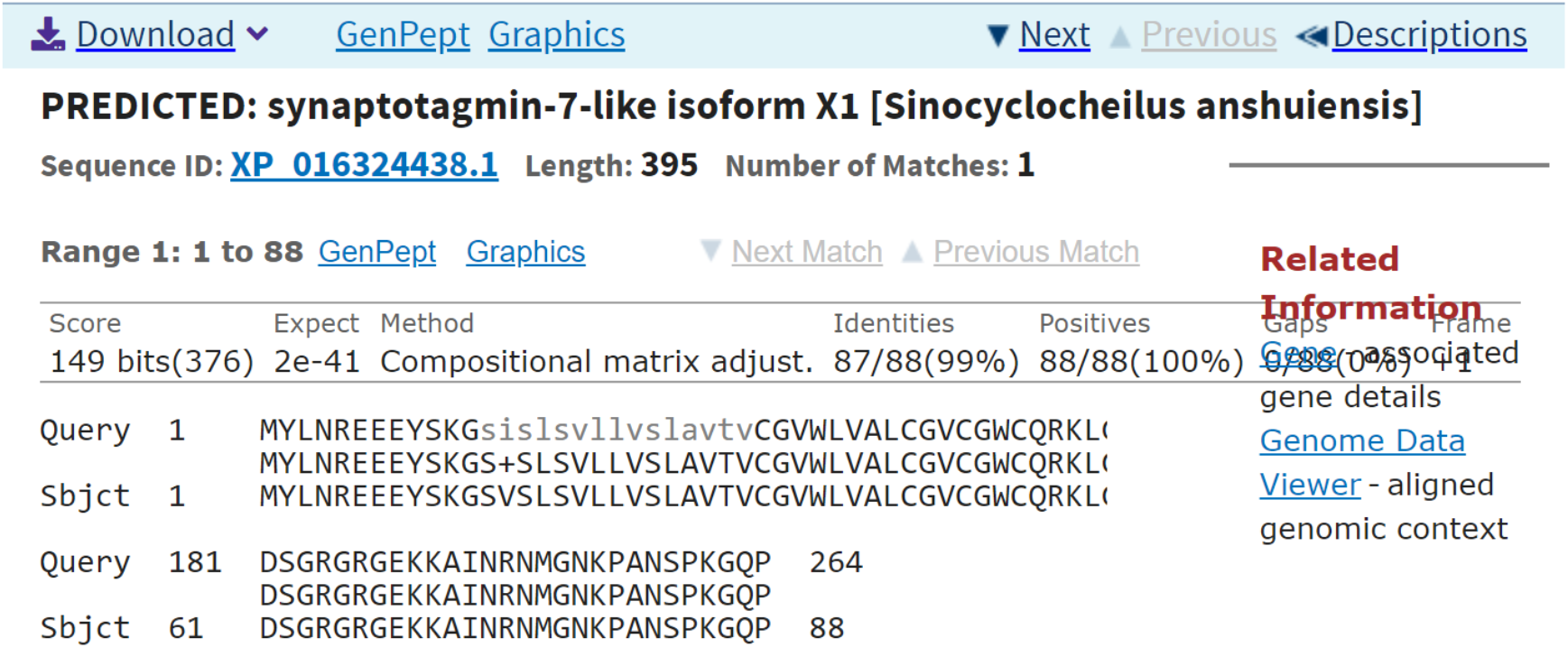

